# *Salmonella* exploits metal-responsive UMPylation to enable intracellular biphasic growth and antibiotic tolerance

**DOI:** 10.64898/2026.02.09.704795

**Authors:** Weiwei Wang, Haihong Jia, Yongyu Wang, Qingjie Bai, Ruirui Liu, Ting Sun, Yingying Yue, Xiaozhi Wang, Yunzhuo Zhao, Xiaoyu Liu, Xilu Yuan, Rongxian Xie, Nannan Song, Yuzhen Wang, Zhongrui Ma, Qixin Chen, Bingqing Li

**Author notes:** These authors contributed equally (Weiwei Wang, Haihong Jia, Yongyu Wang, Qingjie Bai). Correspondence (B.L.), (Q.C.).

## Abstract

*Salmonella* develops profound antibiotic tolerance during *in vivo* infection, directly contributing to antimicrobial treatment failure. Although phenotypic heterogeneity in bacterial growth rates underlies this tolerance, the exact mechanisms remain elusive. Here, we reveal that *Salmonella* exhibits a biphasic growth pattern within macrophages, with initial antibiotic levels modulating lag phase duration. Crucially, this growth plasticity is orchestrated by metal ion homeostasis and a reversible PTM-mediated protein phase transitions. Mechanistically, under stress conditions, enhanced Mn^2+^ uptake activates the UMPylator YdiU, which modifies critical translational machinery components such as elongation factor TufA. These modifications induce protein aggregation, leading to translational arrest and growth stasis. Conversely, upon stress alleviation, *Salmonella* accumulates Mg^2+^ and effluxes Mn^2+^, then the de-UMPylase YdiV is activated, which resolves protein aggregates to enable rapid bacterial resuscitation and proliferation. This metal-responsive proteostatic switch represents an evolutionarily conserved strategy for bacterial stress adaptation, with broad implications for combating antimicrobial tolerance.

**SIGNIFICANCE STATEMENT:** Antibiotic tolerance, a key cause of treatment failure in chronic infections, is often linked to bacterial phenotypic heterogeneity, yet the underlying regulatory mechanisms remain largely understood. This study uncovers a fundamental survival strategy employed by intracellular pathogen *Salmonella*, where host-derived manganese acts as a critical signal to induce a reversible, dormant state. We demonstrate that stress-induced Mn^2+^ influx activates widespread protein UMPylation, which functions as a molecular glue to drive the aggregation of essential proteins, thereby halting bacterial growth and conferring antibiotic tolerance. Crucially, this state is reversible upon stress relief through Mn^2+^ efflux and subsequent de-UMPylation, enabling rapid bacterial resuscitation. As an evolutionarily conserved mechanism of adaptive tolerance, this metal-ion-responsive switch, mediated by reversible protein UMPylation, identifies a key target for developing therapeutics against persistent bacterial infections.

## INTRODUCTION

The persistent global burden of bacterial infections remains a significant public health challenge, with antimicrobial resistance (AMR) contributing to 4.95 million annual deaths^1,2^. Current antibiotic therapies face mounting limitations as resistance mechanisms have been documented across all antimicrobial classes utilized in clinical practice^3^. Notably, pathogens demonstrating *in vitro* susceptibility can develop phenotypic tolerance during host infection, resulting in therapeutic failure, chronic infection persistence, and increased risks of recurrent disease episodes^4,5^.

Antimicrobial treatment failure can primarily be attributed to three bacterial survival strategies: resistance, tolerance, and persistence^4,6^. Resistance refers to acquired microbial capacity to counteract antibiotics, characterized by elevated minimum inhibitory concentrations (MIC)^4^. In contrast, tolerance and persistence involve mechanisms by which bacteria evade the lethal effects of antibiotics, typically through decreased growth rates, resulting in unchanged MIC but significantly prolonged killing times^4,7^. Persister cells represent a small subpopulation (typically<1%), that enters a transient non-replicating state under antibiotic pressure^8,9^. Multiple mechanisms have been implicated in the induction of the persister phenotype^10^, including the activation of the stringent response^11–13^, the involvement of toxin-antitoxin systems^14–17^, alterations in intracellular ATP levels^18–20^ and protein aggregation^21–24^. In contrast, tolerance refers to a population-wide adaptive response characterized by a slowdown in growth kinetics during exposure to antimicrobial agents^25–27^. Current evidence suggests that conventional factors influencing persister cell formation fail to explain pathogen antibiotic tolerance *in vivo*^5,28^. Although preliminary studies have implicated nutritional availability, metabolic activity, and membrane composition and voltage in shaping *in vivo* tolerance, the precise molecular mechanisms remain elusive^5,29–33^.

The phenotypic manifestation of bacterial tolerance fundamentally stems from growth rate modulation within host environments^25–28,34,35^. Upon entering macrophages, *Salmonella* exhibits heterogeneity in survival, with a significant portion entering a dormancy-like state without replication, while another subset remains actively replicative^36–38^. Recent investigations reveal bacterial heterogeneous growth patterns during tissue colonization, characterized by divergent subpopulations exhibiting either proliferative activity or metabolic quiescence^37–41^. While nutrient deprivation could theoretically induce passive growth arrest, emerging data challenge this paradigm by demonstrating transcriptional reprogramming events and a metabolically active state in slowly dividing pathogens^42–44^. These observations suggest potential active regulatory mechanisms governing growth deceleration, though definitive molecular pathways connecting environmental sensing to replication control remain to be elucidated.

In this study, we reveal the mechanisms by which *Salmonella* dynamically regulates its growth within host cells. Under antibiotic exposure, intracellular *Salmonella* follows three distinct fates. A resuscitating subpopulation displays biphasic growth-a lag phase, whose length depends on the initial antibiotic concentration, followed by exponential growth. This adaptive mechanism is mediated through coordinated actions of bacterial metal ion transporters and a unique post-translational modification (PTM) system-UMPylation. Under stress conditions, enhanced Mn^2+^ uptake activates the UMPylator YdiU, which modifies critical translational machinery components such as elongation factor TufA. These modifications induce protein aggregation, leading to translational arrest and growth stasis. Crucially, the enzyme responsible for catalyzing de-UMPylation has been identified as the protein encoded by *ydiV*, which is located directly upstream of the *ydiU* gene. Upon stress alleviation, accumulated Mg^2+^ activates YdiV, which resolves protein aggregates to enable rapid bacterial resuscitation and proliferation. *In vivo* data showed that antibiotic-tolerant Δ*ydiV-U* double mutant *Salmonella* was reduced by five orders of magnitude in host liver compared to the wild-type strain. Taken together, our data showed that *Salmonella* exploits host metal ion gradients coupled with reversible PTM-mediated protein phase transitions to dynamically regulate translation efficiency, establishing an antibiotic tolerance paradigm through real-time growth rate modulation.

## RESULTS

### YdiU governs *Salmonella* antibiotic clearance within the host environment

Our previous studies identified YdiU as a stress-inducible enzyme mediating UMPylation in *Salmonella*, with demonstrated regulatory roles in heat shock response, flagellar synthesis, and iron uptake^45–47^. Notably, YdiU-deficient *Salmonella* exhibited significant enhanced virulence compared to wild-type strains in animal models, yet displayed complete susceptibility to antibiotic treatment - a striking contrast to the antibiotic tolerance typically observed in wild-type *Salmonella* infections (Fig.1a-b). As *Salmonella* is a prototypical intracellular pathogen residing in host macrophages, we established an infection model using murine RAW264.7 macrophages. Intracellular survival rates of WT, Δ*ydiU*, and D248A mutants were quantified within macrophages following supplementation of culture medium with 100×MIC ceftazidime/norfloxacin (Fig.1c-e). At 24 h post-infection, intracellular Δ*ydiU* and D248A mutant were completely eradicated by antibiotic treatment, whereas WT maintained substantial survival (10^3^ CFU/mL). Intriguingly, *in vitro* time-kill assays revealed congruent survival kinetics across three strains under ceftazidime/norfloxacin treatment, establishing YdiU-dependent antibiotic tolerance as an *in vivo*-restricted phenotype (Fig.1f-g). MIC determination showed indistinguishable susceptibility profiles, excluding resistance-mediated clearance defects (Fig.1h). Conclusively, YdiU operates exclusively in host-adapted tolerance without influencing intrinsic resistance or *in vitro* persister development.

**Fig. 1.**
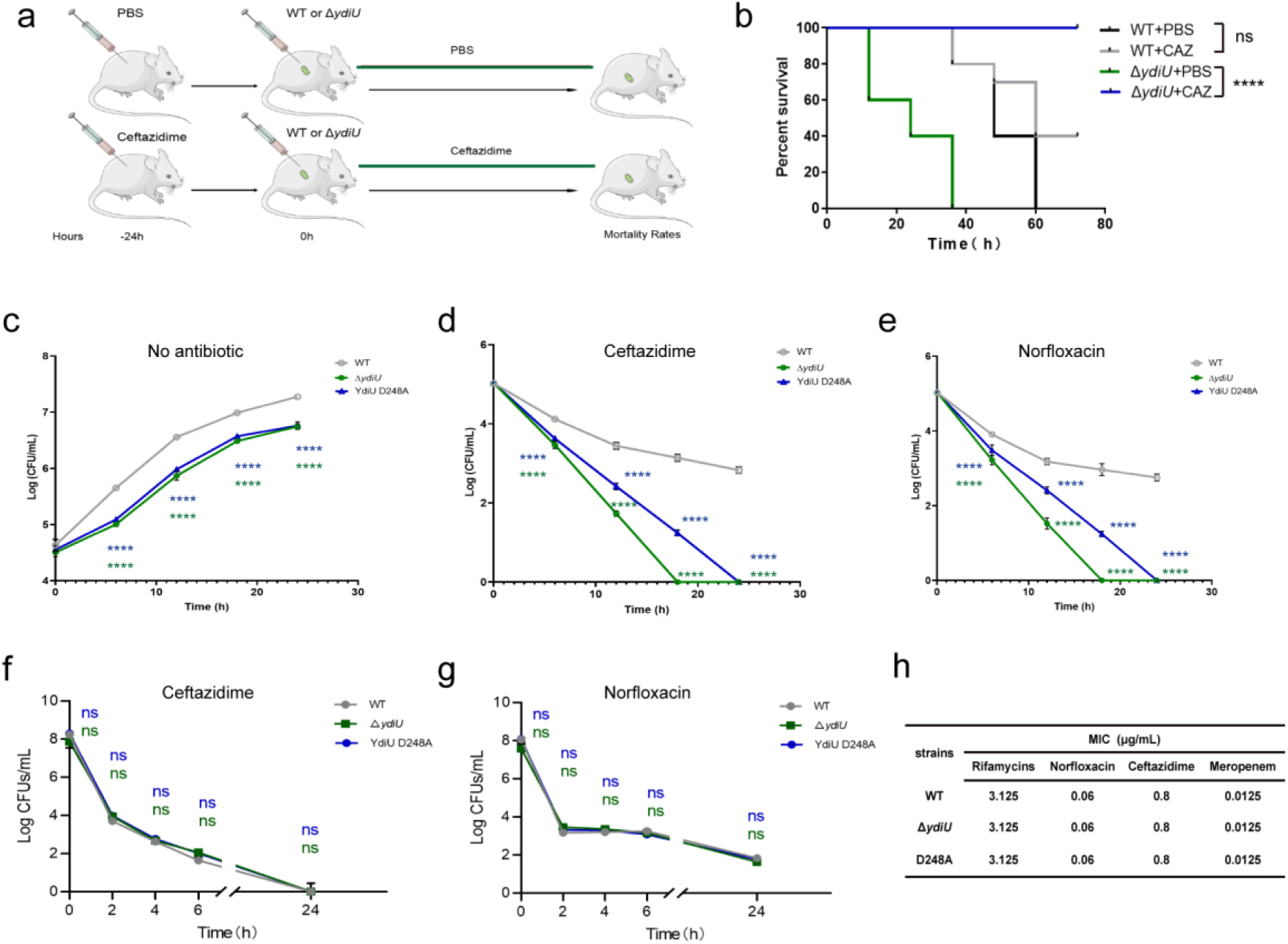
YdiU promotes *Salmonella* antibiotic tolerance within host. (**a**) Antibiotic treatment schematic: Mice pre-treated with antibiotics one day prior to infection, followed by daily post-infection administration (day 0 = infection). (**b**) Survival curves of mice infected with WT or Δ*ydiU Salmonella* under antibiotic treatment (n=5). (**c**–**e**) Intracellular survival of indicated strains in macrophages: Untreated (**c**), or with ceftazidime (**d**) or norfloxacin (**e**). (**f**–**g**) *In vitro* survival of indicated strains after ceftazidime (**f**) or norfloxacin (**g**) exposure. (**h**) Minimum inhibitory concentrations (MICs) against indicated antibiotics. Statistical notes: WT vs. mutant comparisons unless specified; significance denoted as *p < 0.05, **p < 0.01, ***p < 0.001, ****p < 0.0001; ns, not significant.

### YdiU inhibits protein translation and bacterial growth

Broad-spectrum antibiotic screening showed that the Δ*ydiU* strain exhibited up-regulated persister formation under gentamicin treatment (targeting the 30S ribosomal subunit) (Extended Data Fig.1a). Prior work demonstrated that YdiU co-expression significantly suppresses heterologous protein production[45]. Given that numerous toxin-antitoxin systems modulate antibiotic persistence by targeting translation machinery^12,15,48–52^, we hypothesized that YdiU might govern antibiotic tolerance through translational perturbation. To investigate the regulatory role of YdiU in the protein translation process, we developed an *in vitro* translation system (Extended Data Fig.1b). Introduction of YdiU into this system resulted in a significant decrease in protein expression levels (Fig.2 a-b, Extended Data Fig.1c). Furthermore, a real-time expression system for luciferase was established in both WT and Δ*ydiU Salmonella* (Extended Data Fig.1d). The expression levels of luciferase in Δ*ydiU* strain were markedly higher than those observed in WT, demonstrating YdiU-mediated regulation of protein biosynthesis *in vivo* (Fig.2c). To investigate the impact of YdiU on bacterial growth dynamics, we engineered an inducible *Salmonella* strain capable of arabinose-dependent YdiU expression. The growth curve suggested that the induction of YdiU caused marked bacterial growth suppression (Extended Data Fig.1e). Notably, this inhibitory effect consistently manifested during stationary-phase growth, regardless of induction was initiated during logarithmic or stationary phases.

**Fig.2.**
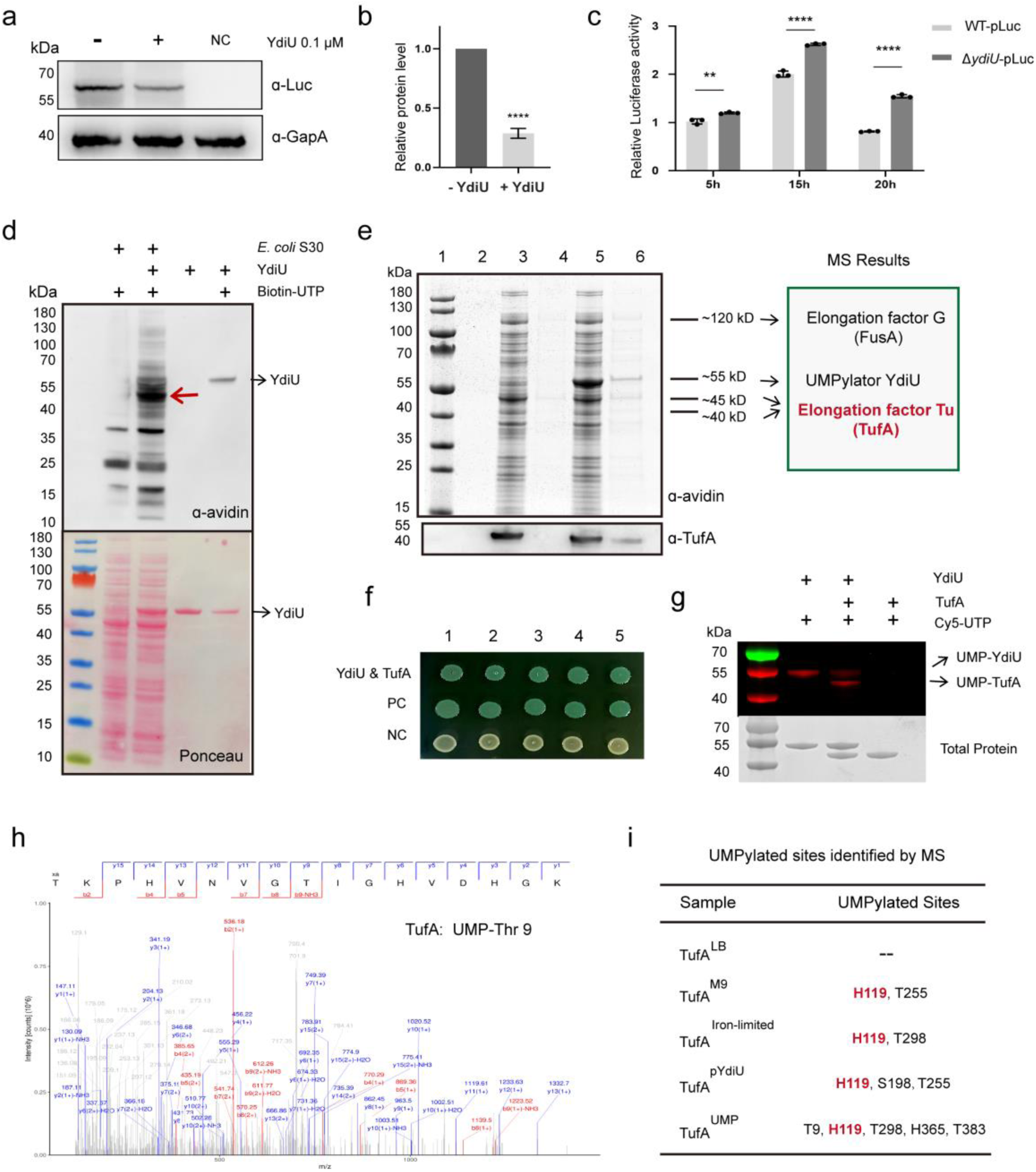
YdiU inhibits protein translation via TufA UMPylation. (**a**) *In vitro* luciferase translation assay ± YdiU. (**b**) Relative luciferase levels (normalized to GapA, densitometric analysis; mean ± SD, n=3). (**c**) Luciferase activity kinetics in WT-pLuc vs. Δ*ydiU*-pLuc strains. (**d**) UMPylation of E. coli S30 extract by YdiU. (**e**) Streptavidin pulldown with biotinylated UTP: SDS-PAGE (Lane 4: -YdiU control; Lane 6: +YdiU). Target bands identified by mass spectrometry (MS), validated by TufA immunoblotting. (**f**) Bacterial adenylate cyclase two-hybrid assay: YdiU-TufA interaction (PC: positive control; NC: negative control). (**g**) *In vitro* UMPylation of purified TufA by YdiU using Cy5-UTP. (**h**) Representative MS/MS spectra of UMPylated TufA peptides. (**i**) Evolutionary conservation of identified UMPylation sites in TufA homologs. *Statistical significance: **p < 0.01; ***p < 0.0001 (b, c).

### YdiU UMPylates elongation factor TufA of protein translation system

We previously identified YdiU as an enzyme that catalyzes UMPylation of proteins^45^. Therefore, we propose that YdiU regulates translation by UMPylating specific protein(s) in the *in vitro* translation system. Then, we conducted UMPylation experiments utilizing total proteins from the translation system as the target. Intriguingly, YdiU UMPylates multiple proteins within the translation system (Fig.2d). The predominant modified band falls within the 40-50 kDa range (Fig.2d). To further identify the above UMPylated proteins, a screening system based on the streptavidin-biotin interaction was established (Extended Data Fig.1f). Proteomic analysis of the enriched fractions revealed two predominant Coomassie-stained bands migrating at 55 kDa and 45 kDa, which were assigned to YdiU and elongation factor Tu (TufA) through mass spectrometry (Fig.2e). Western blot analysis confirmed TufA presence specifically in YdiU-modified samples (Fig.2e).

TufA is one of the most abundant proteins in bacterial cells and plays a crucial role in translation. Additionally, many toxin specifically target TufA^50,53–55^. Then, we hypothesize that YdiU may inhibit translation by modulating TufA. A bacterial two-hybrid experiment confirmed the strong interaction between YdiU and TufA *in vivo* (Fig.2f). Furthermore, purified YdiU and TufA were conducted one-on-one UMPylation experiments *in vitro*. YdiU is capable of directly UMPylating TufA (Fig.2g). To further identify the UMPylated sites of TufA by YdiU, TufA proteins from LB medium (TufA^LB^), M9 medium (TufA^M9^), Iron-limited medium (TufA^iron-limited^), co-expressed with YdiU (TufA^pYdiU^) and TufA *in vitro* UMPylated by YdiU (TufA^UMP^) were prepared for mass spectrometry analysis. A total of seven UMPylated sites were detected, distributing across three structural domains of TufA (Fig.2h-i and Extended Data Fig.2a). Among these, H119 was identified as a site present in both *in vivo* and *in vitro* modified samples. H119 located in the GTP binding pocket of TufA indicating that UMPylation on H119 may directly influence GTP binding (Extended Data Fig.2b). The residues surrounding the modification sites exhibited the previously observed characteristics of YdiU-catalyzed UMPylated sites, the predominance of hydrophobic and charged residues (Extended Data Fig.2c).

### UMPylated TufA undergoes phase separation both *in vitro* and *in vivo*

UMPylation of TufA caused reaction solution turbidity (Extended Data Fig.3a), unlike the insoluble aggregates previously observed in UMPylated GroEL or Fur ^45,46^. To investigate UMPylation-driven phase separation, GFP-TufA constructs were engineered. Phase-separated condensates formed exclusively in UMPylated TufA reactions, whereas unmodified TufA showed no condensation under identical conditions (Fig.3a and Extended Data Fig.4a-b). Additionally, photobleaching experiments confirmed that phase separation of UMPylated TufA is dynamic (Fig.3b-c). Fluorescent colocalization analysis with Cy3-UTP demonstrated UMPylation-induced LLPS of TufA, as evidenced by precise overlap between Cy3 signals and condensates (Extended Data Fig.4c). Kinetic monitoring revealed temporal progression from metastable LLPS (30 min) to irreversible aggregation (>60 min) (Extended Data Fig.4d). Notably, TufA lacks intrinsic disordered regions (Extended Data Fig.2d). Biophysical analysis showed that UMPylation induces secondary structural perturbations (Extended Data Fig.3b-c) and increases the hydrodynamic radius (from 2.7 nm to >34.2 nm; Extended Data Fig.3d-f), indicating that UMPylation-mediated aggregation results from secondary structural changes.

**Fig.3.**
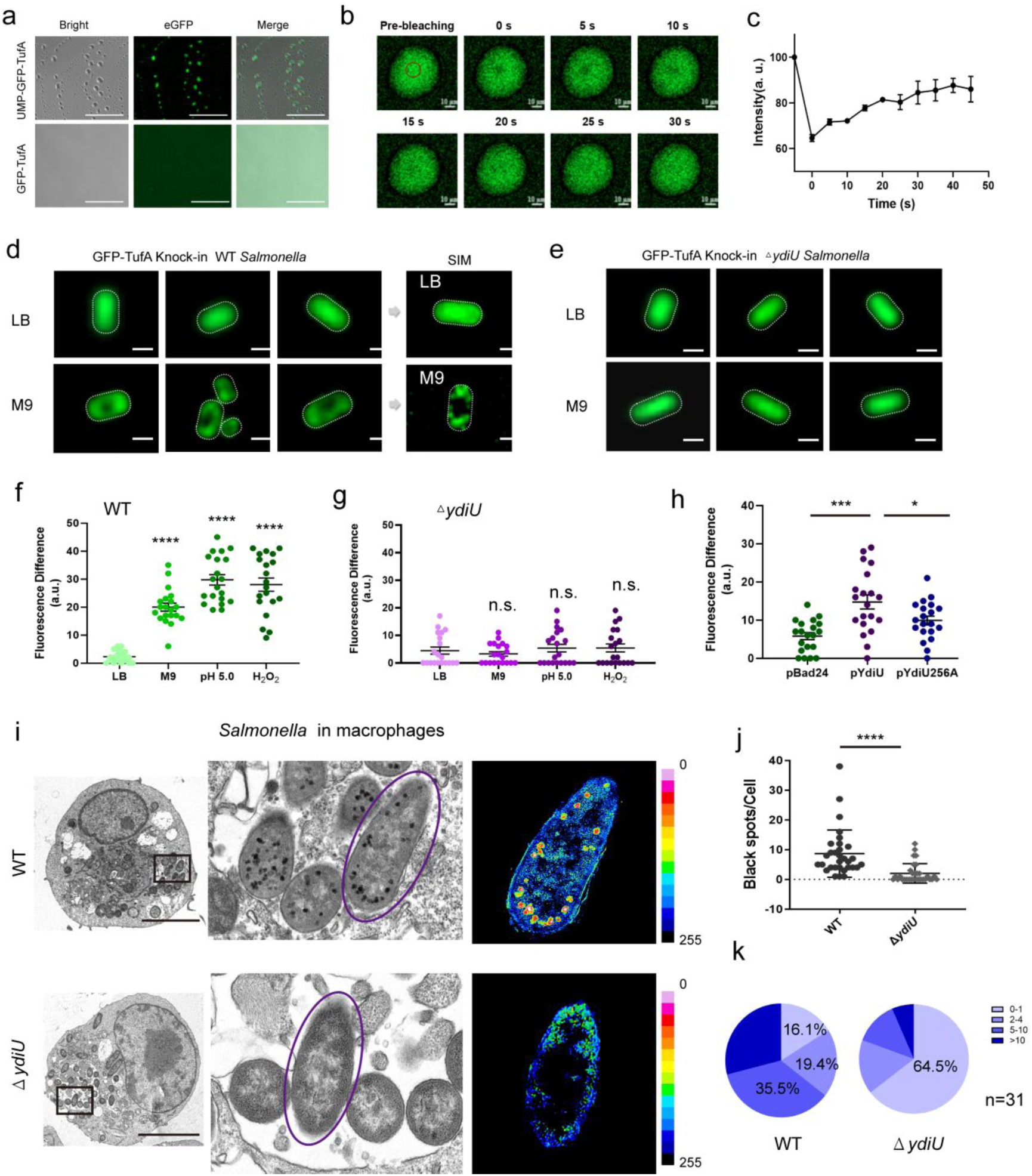
TufA undergoes UMPylation-mediated phase separation across *in vitro*, bacterial, and infected cellular environments. (**a**) *In vitro* phase separation of GFP-TufA ± Mn²⁺ (UMPylation cofactor). Scale bar = 50 μm. (**b**) FRAP analysis of GFP-TufA condensates. Scale bar = 10 μm. (**c**) FRAP recovery kinetics (mean ± SD; n = 3). (**d**-**e**) Subcellular localization of genomically integrated GFP-TufA in WT (**d**) vs. *ΔydiU* (**e**) *Salmonella* under nutrient-rich (LB) or minimal (M9) conditions. Scale bar = 1 μm. (**f**-**g**) Quantification of intracellular GFP-TufA fluorescence intensity in WT (f) and *ΔydiU* (g) strains (n = 20 cells/condition). (**h**) Complementation assay: GFP-TufA intensity in WT harboring pBAD24 (vector), pBAD24-YdiU, or pBAD24-YdiU D248A ± L-arabinose (n = 20). (**i**) TEM of infected RAW264.7 macrophages showing electron-dense aggregates in WT vs. *ΔydiU Salmonella* at 10 hpi. Scale bar = 5 μm. (**j**) Quantification of bacterial aggregates per infected macrophage (10 hpi). (**k**) Distribution of intracellular bacterial foci per macrophage infected with WT vs. *ΔydiU* strains (10 hpi). Significance significance: *p < 0.05, ***p < 0.001, ****p < 0.0001; ns, not significant.

To investigate the *in vivo* phase separation behavior of TufA, we generated strains overexpressing GFP-tagged TufA. Fluorescence imaging revealed robust formation of TufA phase-separated condensates in WT cells cultured in YdiU-inducing medium. In contrast, Δ*ydiU* strains exhibited no detectable TufA phase separation (Extended Data Fig.3g-h). FRAP assays revealed <15% signal restitution in TufA condensates from YdiU-expressing cells, demonstrating UMPylation drives irreversible aggregation rather than dynamic phase separation *in vivo*. To eliminate the possibility of false positives caused by over-expression, we constructed strain of WT and Δ*ydiU Salmonella* with a GFP inserted in situ at the *tufA* gene (GFP-TufA knock-in). Under YdiU-inducing conditions, TufA forms condensates in wild-type *Salmonella*, whereas Δ*ydiU* mutants lack this phenotype (Fig.3d-g). Using the TufA knock-in strain, we generated YdiU or catalytically inactive YdiU D256A overexpressed strains (pYdiU/pYdiU256A). Strikingly, YdiU triggered TufA spatial redistribution under non-stress conditions, dependent on its UMPylation activity (Fig.3h and Extended Data Fig.3i).

### YdiU-mediated UMPylation acts as a common trigger to drive phase separation

Additional YdiU substrates (e.g., GroEL, Fur) exhibit UMPylation-dependent aggregation, all lacking intrinsically disordered regions (IDRs)^45,46^. Then we propose UMPylation may serve as a universal driver of protein phase separation. To test this hypothesis, we generated fluorescently tagged target proteins and revealed that GroEL, Fur, and FlhD are all capable of undergoing phase separation following UMPylation (Extended Data Fig.5a-c). Notably, mirroring TufA behavior, LLPS initiates promptly post-UMPylation, with progressive modification driving condensate formation that ultimately transitions to irreversible precipitation. Building on these findings, we propose that YdiU may induce widespread UMPylation of bacterial proteins, promoting their aggregation. To test this hypothesis, isogenic YdiU-expressing strains were constructed in the Δ*ydiU* background. Transmission electron microscope (TEM) analysis of intracellular ultrastructure revealed striking differences: 76% of YdiU-expressing cells harbored substantial melano-aggregates, versus only 11% in Δ*ydiU* controls (Extended Data Fig.5d-e), establishing YdiU-mediated UMPylation as a key driver of protein aggregation both *in vivo* and *in vitro*.

### *Salmonella* undergoes YdiU-dependent protein aggregation within macrophage

Upon macrophage invasion, *Salmonella* markedly upregulates YdiU expression^46,47^, then the protein aggregation states of *Salmonella* within macrophage were determined by TEM. TEM time-course analysis of wild-type *Salmonella*-infected macrophages revealed dynamic melano-aggregate formation: characteristic dark proteinaceous foci emerged in bacterial cells at early stage during infection (2-10 hpi), which dynamically disassembled during late infection stages and almost completely vanished prior to host cell lysis (Extended Data Fig.5f-g). Notably, Δ*ydiU* mutants displayed profound defects in melano-aggregate biogenesis, with the majority of Δ*ydiU* completely lacking these structures versus wild-type (WT) controls (Fig.3i-j). Quantitative analysis at 10 hpi revealed stark disparities: 64.6% of WT bacteria harbored ≥5 melano-aggregates/cell, contrasting with merely 19.3% in Δ*ydiU* populations (Fig.3k), establishing YdiU as the master regulator of transient biomolecular condensation in intracellular *Salmonella*. Critically, the YdiU D248A mutant strain failed to form protein aggregates at any time point during macrophage residence, definitively demonstrating that YdiU-mediated UMPylation activity is essential for protein aggregate formation in intracellular *Salmonella* (Extended Data Fig.5h).

### YdiV acts as a Mg^2+^/Ca^2+^-dependent de-UMPylator

*Salmonella* surviving macrophage infection develops YdiU-dependent electron-dense aggregates. However, the time-dependent disassembly of these biomolecular condensates implies that bacterial de-UMPylated enzymes may actively dismantle these structures during late infection stages. Given that toxins and antitoxins are typically located in close proximity within the genome, we examined the genes adjacent to YdiU. Notably, we identified YdiV, the situated upstream of the YdiU gene, as particularly significant (Fig.4a, Extended Data Fig.6a). YdiV belongs to the EAL-like protein family, which primarily functions to hydrolyze the signaling molecule c-di-GMP^56^. In YdiV, half of the amino acids responsible for binding and hydrolyzing c-di-GMP are mutated and have been confirmed to lack the capability to degrade c-di-GMP^57^. Given that the nucleotide-binding pocket in YdiV is still present^58^, we speculate whether it could catalyze de-UMPylation. To investigate this, we developed a de-UMPylation system utilizing UMPylated YdiU or UMPylated TufA as the substrates (Extended Data Fig.7a). Our findings demonstrate that YdiV possesses de-UMPylation activity, which is activated by both Ca²⁺ and Mg²⁺ ions (Fig. 4b). Notably, YdiV can catalyze the removal of UMP moieties from both UMP-YdiU and UMP-TufA (Fig.4c-d, Extended Data Fig.7b-c). To determine the molecular mechanism by which YdiV catalyzes de-UMPylation, molecular dynamics simulations of YdiV&UMP-Tyr&Ca^2+^ complex were performed (Fig.4e). Ca^2+^ were located into the complex structure similar with that observed in other EAL homologous proteins (Extended Data Fig.6b). Glu29 and Glu116 are involved in Ca^2+^-binding and YdiV-UMP interaction (Fig.4e), and these amino acids are highly conserved within the YdiV protein family (Extended Data Fig.6c). Structural analyses reveal YdiV’s deUMPylation mechanism operates through a three-step cascade: Glu116-activated water generates a hydroxyl nucleophile for stereospecific ɑ-phosphate attack, forming a pentacoordinate phosphorane transition state, followed by coordinated bond cleavage releasing UMP and restoring tyrosine residues (Extended Data Fig.6d). The mutation of Glu116 deprives de-UMPylation activity of YdiV, supporting this mechanistic hypothesis (Fig.4f-g). In conclusion, YdiV functions as a de-UMPylator, representing a novel phosphotransferase subfamily diverging from classical EAL fold biochemistry.

**Fig.4.**
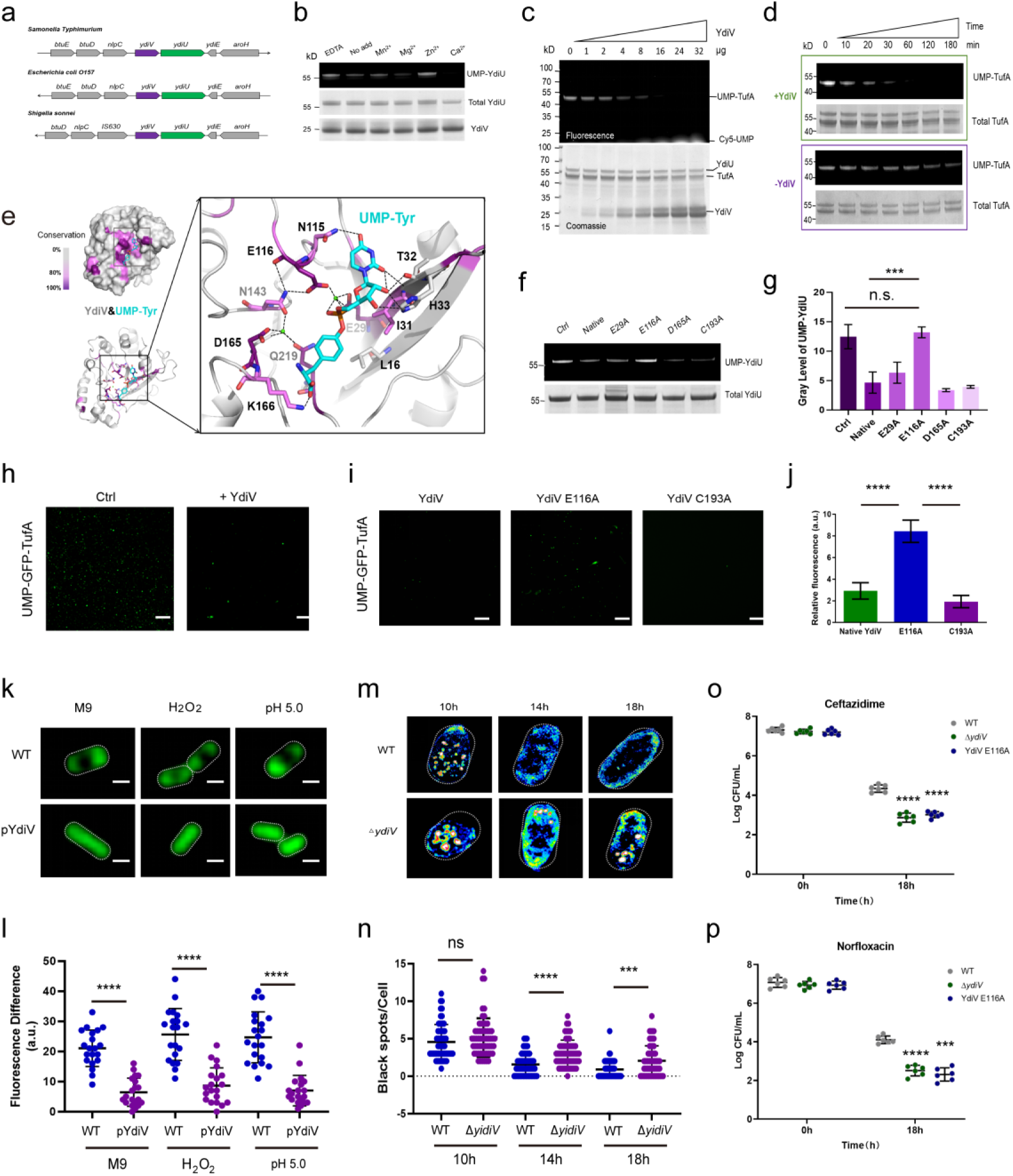
YdiV catalyzes de-UMPylation and modulates protein aggregation and bacterial tolerance. (**a**) Genomic colocalization of *ydiV* and *ydiU* in pathogenic bacteria. (**b**) *In vitro* deUMPylation of UMP-YdiU by YdiV with indicated metal cofactors/EDTA. (**c**) Concentration-dependent deUMPylation by YdiV (60 min; UMP-TufA detected by autoradiography). (**d**) Time-course of deUMPylation by YdiV. (**e**) Molecular dynamics structure of YdiV-UMP-Tyr-Ca²⁺ complex (conserved residues in purple). (**f**) DeUMPylation activity of WT YdiV versus point mutants (1 h reaction). (**g**) Densitometric quantification of autoradiography bands in (j) (n=3). (**h**-**i**) *In vitro* phase separation of GFP-TufA ± 10 µg WT YdiV (H) or mutants (i) (scale bars: 50 µm). (**j**) Aggresome quantification from (h-i) (n=20). (**k**) Subcellular distribution of genomic GFP-TufA in WT vs. pYdiV Salmonella under stress (scale bar: 1 µm). (**l**) Fluorescence intensity quantification for (k) (n=20 cells). (**m**) Representative TEM images of intracellular aggregates in WT vs. pYdiV Salmonella-infected RAW264.7 macrophages. (**n**) Quantification of aggregates per cell in (m) (n=30-50 cells). (**o**-**p**) Intracellular survival in antibiotic-treated macrophages: ceftazidime (o), norfloxacin (p). Significance significance: ***p < 0.001, ****p < 0.0001; ns, not significant.

### YdiV modulates protein disaggregation and antibiotic tolerance during infection

To investigated the role of YdiV in UMPylation-mediated protein aggregation, YdiV and E116A mutant were introduced into the *in vitro* phase separation system of UMPylated TufA. The native YdiV significantly suppresses phase separation phenomenon of GFP-TufA, while YdiV E116A mutant does not (Fig.4h-j). To delineate YdiV’s role in stress-induced TufA phase separation in bacterial cells, we engineered a *Salmonella* strain with constitutive YdiV expression (pYdiV). Strikingly, under equivalent stress regimens, pYdiV cells exhibited homogeneous TufA distribution versus pronounced uneven distribution in wild-type controls (Fig.4k-l), establishing YdiV’s capacity to reverse stress-triggered protein aggregation *in vivo*.

Then, the formation of intracellular protein aggregates in Δ*ydiV* during macrophage infection was determined. In the early infection stage, Δ*ydiV* formed protein aggregates at levels comparable to WT. However, as infection progressed, 95% of WT bacteria cleared these aggregates, whereas over one-third of Δ*ydiV* retained abundant aggregates upon host cell lysis (Fig.4m-n). The E116A mutant exhibited a similar aggregate clearance defect as Δ*ydiV* (Extended Data Fig.8). Furthermore, both Δ*ydiV* and E116A mutant strains showed significantly reduced tolerance compared to WT, further confirming that YdiV-dependent de-UMPylation promotes bacterial antibiotic tolerance (Fig.4o-p).

To further characterize the composition of the black aggregates, wild-type *Salmonella,* Δ*ydiV* and pYdiU were cultured and protein aggregate formation was observed using TEM (Extended Data Fig.9a). The results revealed distinct protein aggregates in Δ*ydiV* and pYdiU, but not in WT strain. Then, the aggregates were isolated via density gradient centrifugation and analyzed by SDS-PAGE^20^. Consistent with the TEM findings, substantial aggregated proteins were successfully isolated only from Δ*ydiV* and pYdiU strains (Extended Data Fig.9b). Mass spectrometry analysis identified 311 major protein components in YdiU-induced aggregates and 294 proteins in the Δ*ydiV* strain while only five protein components identified in WT control (PSM>10) (Extended Data Fig.9c-d). Protein clustering analysis revealed that the aggregates were predominantly composed of components of the protein translation machinery (including elongation factors, ribosomal proteins, and aminoacyl-tRNA synthetases), molecular chaperones, and RNA polymerase, and metabolic enzymes (Extended Data Fig.9e-i). Notably, the vast majority of proteins (42/46) previously identified in UMPylated proteome profiling upon YdiU expression^45^ were detected in the insoluble fraction, although most of them are classified as soluble and not known to undergo phase separation (Extended Data Fig.9d-i). This results provide further evidence supporting the hypothesis that YdiU-mediated UMPylation induces widespread protein aggregation.

### YdiU-YdiV facilitates intracellular biphasic growth to evade antibiotic clearance

Above data reveals that the YdiU-YdiV system mediates reversible protein aggregation during *Salmonella* survival in macrophages. Given established links between protein aggregation and bacterial dormancy^20,59^ and the fact that YdiU inhibits protein translation and growth rate, we hypothesize YdiU-YdiV system might modulate host-adapted growth states of *Salmoenlla*. Live single-cell imaging of intracellular wild-type *Salmonella* exposed to antibiotics revealed the existence of three phenotypically distinct subpopulations within macrophages (Fig.5a-b): Rapidly cleared (186/359 in 25*MIC group), Tolerant replicators (151/359; 87.3% of survivors) and Non-replicating persisters (22/359; 12.7%). Notably, tolerant replicators exhibited biphasic growth kinetics with extended adaptation phases (lag time) proportional to antibiotic concentration (Fig.5c). Strikingly, Δ*ydiU* mutant showed minimal lag phases and robust early replication, while Δ*ydiV* strains failed to initiate rapid proliferation observed in WT (Fig.5d-f). Compared to wild-type strains, the majority of Δ*ydiU* exhibit rapid proliferation in the early stage, while being gradually cleared during the mid-to-late stages (Fig.5e-f). Then we propose a regulatory dichotomy: YdiU enforces early dormancy through UMPylation-dependent translational inhibition, while YdiV licenses subsequent resuscitation through de-UMPylation that resolves aggregates to restart growth.

**Fig.5.**
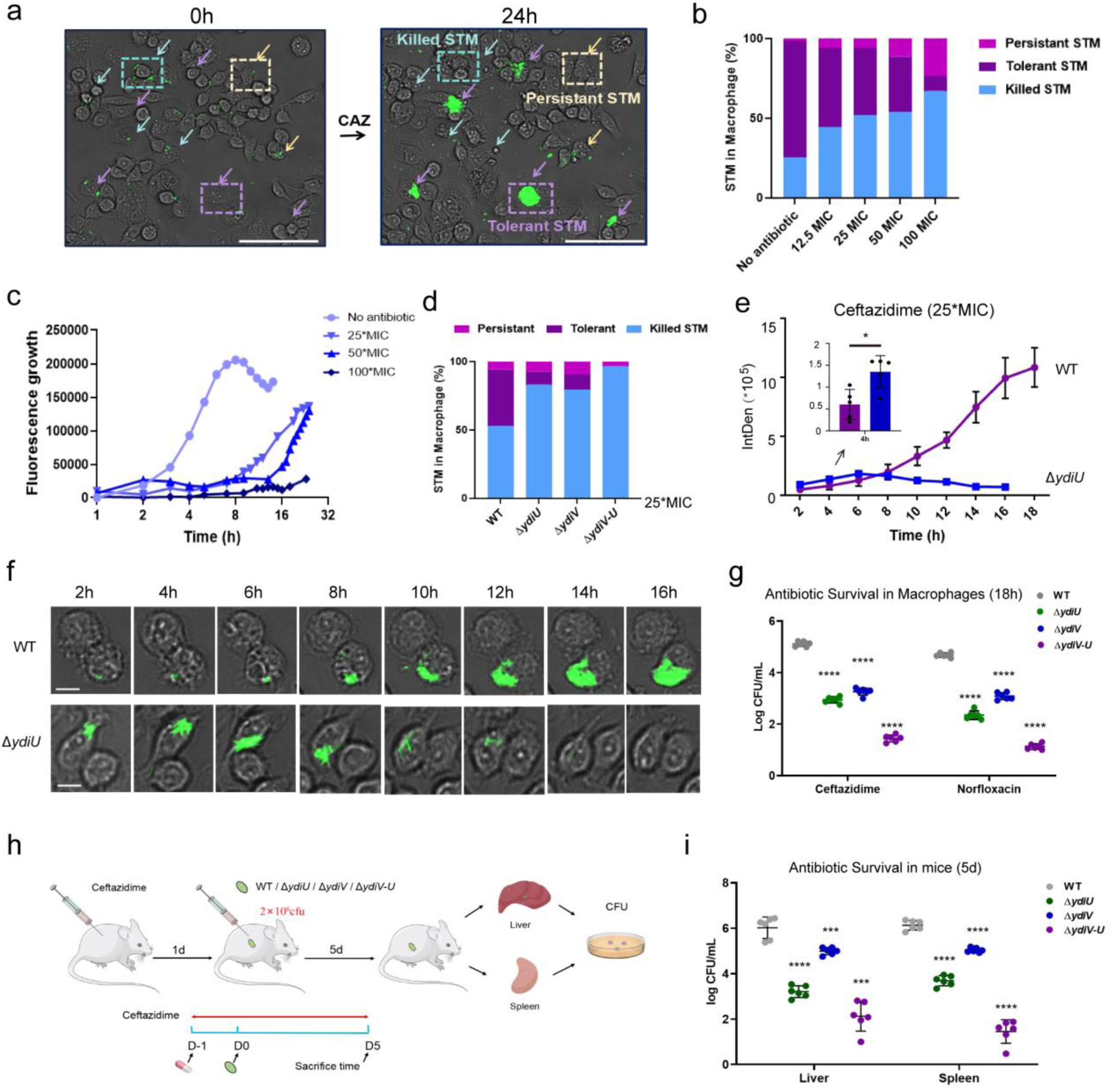
YdiU-YdiV facilitates intracellular biphasic growth to evade antibiotic clearance. (**a**) Intracellular imaging of wild-type *Salmonella* within macrophages before and after 24-hour ceftazidime treatment. Bacteria are labeled with green fluorescence. Blue arrows and dashed borders denote killed *Salmonella*, yellow arrows and borders indicate persisters, and purple arrows and borders mark tolerant *Salmonella*. Scale bar: 100 μm. (**b**) Proportions of WT *Salmonella* in three survival states under varying antibiotic concentrations. 50–100 cells were analyzed per group. (**c**) Proliferation curves of antibiotic-tolerant wild-type *Salmonella* in macrophages under different antibiotic concentrations. (**d**) Proportions of four *Salmonella* strains in three survival states under 25*MIC ceftazidime treatment. 50–100 cells were analyzed per group. (**e**) Fluorescence quantification (reflecting bacterial proliferation) of intracellularly tolerant wild-type Salmonella and Δ*ydiU* mutants post-infection (n=5). (**f**) Representative live-cell images of intracellularly tolerant wild-type *Salmonella* and Δ*ydiU* mutants post-infection. Scale bar: 10 μm. (**g**) Comparative survival of four *Salmonella* strains in macrophages under antibiotic treatment. (**h**) Schematic of the animal experimental workflow. (**i**) Bacterial loads of four strains in murine livers and spleens after 5-day antibiotic therapy (n=6). Significance significance: *p<0.05, ***p < 0.001, ****p < 0.0001.

To assess contributions of YdiU and YdiV to antibiotic tolerance, we quantified intracellular survival of WT, Δ*ydiU*, Δ*ydiV*, and Δ*ydiV-U* strains under ceftazidime/norfloxacin treatment. The Δ*ydiV-U* double knockout exhibited a severe persistence defect (2-3 log reduction in bacterial counts vs. WT), while single knockouts showed intermediate attenuation (1-2 log reduction) (Fig.5g), confirming synergistic roles of YdiV and YdiU in macrophage persistence. These findings were validated *in vivo* using a ceftazidime-treated murine model (Fig.5h). All mutant strains displayed significantly impaired organ colonization compared to WT (Fig.5i-j). Notably, Δ*ydiV-U* infection resulted in near-complete clearance from host tissues (5-log reduction in liver, 4-log in spleen), demonstrating that the YdiU-YdiV axis is indispensable for bacterial antibiotic tolerance during clinical antibiotic therapy.

### *Salmonella* employs Mn^2+^ homeostasis to potentiate antibiotic tolerance

While the YdiU-YdiV axis is critical for *Salmonella*’s antibiotic tolerance in the intramacrophage niche, a key question remains unresolved: How do these two opposing enzymatic systems -YdiU (UMPylase) and YdiV (de-UMPylase)-coordinate temporally distinct roles despite constitutive co-expression from the same operon? Because the expression of both YdiU and YdiV remain high level during *Salmonella* infection^46,60^, the dormancy-revival cycle mediated by YdiU/YdiV is clearly not regulated at the level of protein expression. Given that both proteins function as enzymes, we propose that *Salmonella* achieves dynamic control by modulating their enzymatic activities. Crucially, YdiU and YdiV exhibit distinct metal ion dependencies: YdiU is a Mn²⁺-dependent UMPylator, whereas YdiV requires Mg²⁺/Ca²⁺ for de-UMPylation. This divergence suggests that host-derived fluctuations in metal ion availability may act as biochemical switches to sequentially activate YdiU-mediated growth arrest and YdiV-driven resuscitation. To prove above hypothesis, we systematically monitored the dynamic changes of four metal ions in macrophage during *Salmonella* infection. The levels of Mg^2+^, Zn^2+^, and Ca^2+^ were analyzed using probe-based imaging, while Mn levels were quantified via ICP-MS due to the lack of specific imaging probes (Fig.6a-d). Notably, Mn^2+^ levels were significantly up-regulated in the host cytoplasm 4-10 hours post-infection, while Mg^2+^ accumulated predominantly within bacterial cells during later infection stages (Fig.6a-b). Zn^2+^ concentrations remained stable throughout the process, while Ca^2+^ levels exhibited a pronounced decline post-infection (Fig.6c-d). Strikingly, *in vitro* modeling of antibiotic stress (exposure) versus recovery (removal) uncovered counteractive Mg^2+^/Mn^2+^ regulation in *Salmonella*. Antibiotic challenge drove preferential Mn^2+^ accumulation coupled with Mg^2+^ depletion, while stress termination reversed this ion flux polarity (Fig.6e-h). Then, the impact of environmental metal ions on antibiotic tolerance was detected. Supplementing bacterial cultures with exogenous metal minimally impacts persister cell formation. Conversely, Mn^2+^ addition to macrophage cultures significantly enhances antibiotic tolerance (10- to 30-fold), while EDTA supplementation in macrophage media markedly reduces intracellular survival (8- to 15-fold reduction) (Fig.6i). *Salmonella* primarily acquires Mn^2+^ via the MntH importer, with efflux mediated by MntP and FieF (also known as YiiP)^61,62^ (Fig.6j). Three Mg^2+^ transporters mutants Δ*corA*, Δ*mgtA* and Δ*mgtB* exhibited modest but significant Mg^2+^ depletion (∼50% reduction vs WT) at 12h post-infection, revealing their role in late-stage Mg^2+^ accumulation (Fig.6k). Intracellular tolerance assays identified the Mn^2+^ importer MntH and efflux pumps MntP/FieF as critical contributors to bacterial persistence, whereas knockout of Mg^2+^ transporters (CorA, MgtA, MgtB) or the Zn^2+^ uptake protein ZntB showed minimal effect on intracellular survival (Fig.6l-m). Critically, these mutants minimally affected *in vitro* persister formation, demonstrating that the role in antibiotic tolerance of Mn^2+^ homeostasis are strictly intracellular phenomena. To verify whether metal ion involves in protein aggregation, we employed TEM to track dark aggregate dynamics of intracellular mutant strains. The results demonstrated that MntP facilitates aggregate formation (10 hpi), whereas MntP/FieF promote aggregate disassembly in the late infection stage (Fig. 6p-q). However, the Δ*mgtA* or Δ*mgtB* strain showed no difference from wild-type *Salmonella* in forming intramacrophage aggregates, likely due to failed suppression of late-infection Mg²⁺ accumulation. Collectively, above data demonstrate that Mn^2+^ uptake and efflux contribute to *Salmonella* antibiotic tolerance through protein aggregation-disaggregation dynamics.

**Fig.6.**
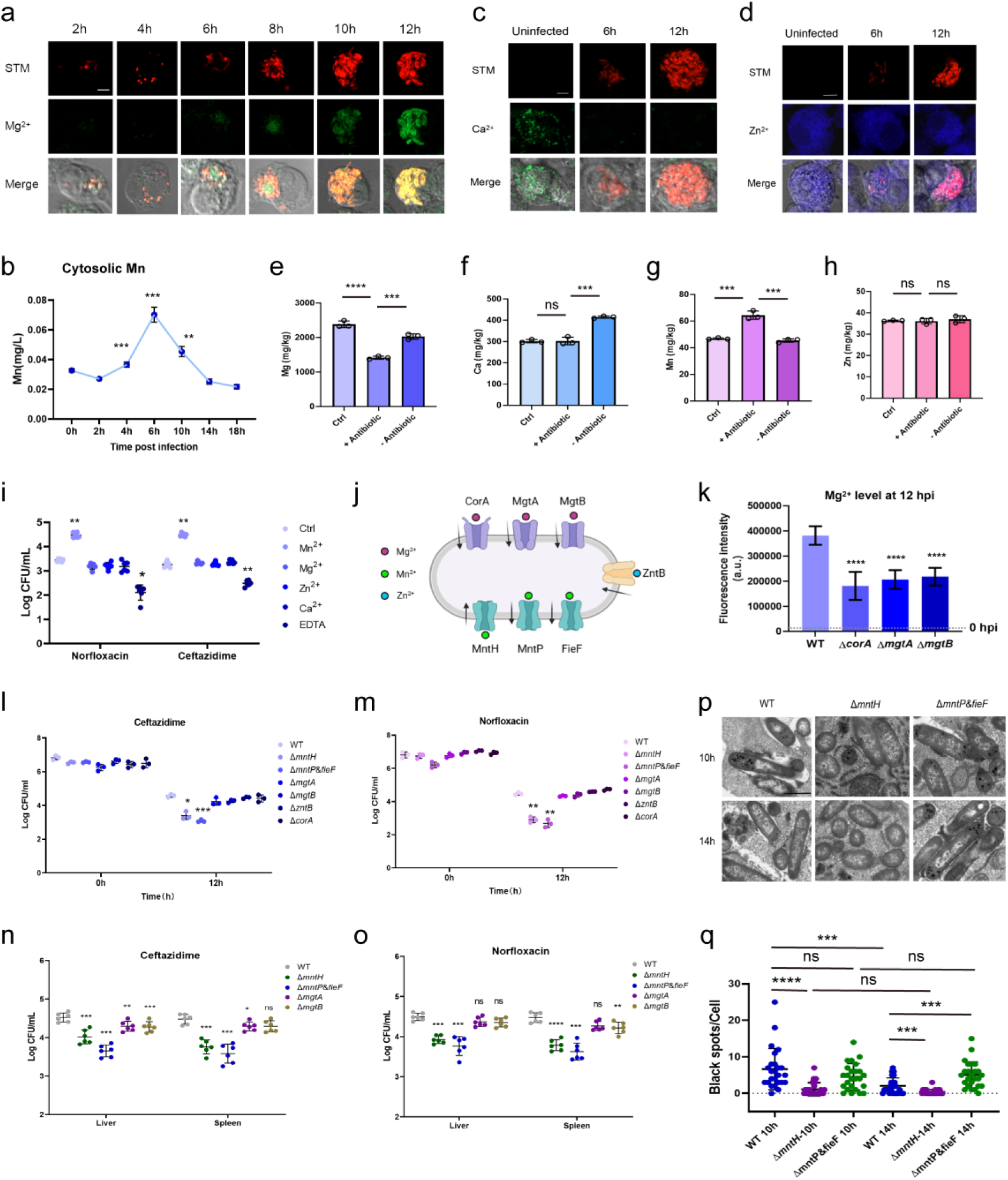
Salmonella employs Mn^2+^ homeostasis to potentiate antibiotic tolerance. (**a**) Intracellular Mg²⁺ distribution in macrophages infected with wild-type *Salmonella* (2–12 h post-infection). Bacteria (red, cherry fluorescence) and Mg²⁺ (green). Scale bar: 2 μm. (**b**) Intracellular Ca²⁺ distribution post-infection with wild-type *Salmonella*. Bacteria (red) and Ca²⁺ (green). Scale bar: 2 μm. (**c**) Intracellular Zn²⁺ distribution post-infection with wild-type *Salmonella*. Bacteria (red) and Zn²⁺ (blue). Scale bar: 2 μm. (**d**) Cytosolic free Mn²⁺ levels in macrophages at different timepoints after wild-type *Salmonella* infection (n=3). (**e**–**h**) Intracellular metal ion content in macrophages infected with wild-type *Salmonella* under antibiotic treatment and post-antibiotic removal: Mg²⁺ (e), Ca²⁺ (f), Mn²⁺ (g), Zn²⁺ (h). (**i**) Survival of intracellular *Salmonella* under ceftazidime/norfloxacin treatment following metal ion supplementation or EDTA chelation in macrophage media. (**j**) Schematic of indicated metal ion transporters on the *Salmonella* membrane. (**k**) Mg²⁺ uptake in macrophages infected with Δ*corA*, Δ*mgtA*, or Δ*mgtB* mutants versus wild-type *Salmonella* at 12 h post-infection. (**l**, **m**) Intracellular survival of metal transporter knockout strains under ceftazidime (l) or norfloxacin (m) treatment at 0 h and 12 h. (**n**, **o**) Bacterial loads of metal transporter knockout strains in murine livers/spleens after 5-day ceftazidime (n) or norfloxacin (o) therapy. (**p**, **q**) Representative image (p) and quantification (q) of black aggregation of Mn²⁺ transporter knockout strains at mid-stage and late-stage infection within macrophages. Significance significance: *p < 0.05, **p < 0.01, ***p < 0.001, ****p < 0.0001; ns, not significant.

### Restriction of host-derived Mn^2+^ effectively promotes *Salmonella* antibiotic clearance

Given the critical role of metal signaling in *Salmonella* antibiotic tolerance, we investigated whether restricting host manganese (Mn) or magnesium (Mg) uptake would impact bacterial antibiotic tolerance. To test this, mice were fed either Mn-restricted or Mg-restricted diets, followed by infection with wild-type *Salmonella*. Results showed that compared to normal diet controls, Mn restriction significantly enhanced antibiotic-mediated bacterial clearance. In contrast, Mg restriction exhibited negligible effects (Fig.7a-b). Subsequent infection experiments using the Δ*ydiV-U* double-knockout strain revealed no enhancement of antibiotic clearance under Mn-restricted conditions. These findings further demonstrate that Mn²⁺ signaling promotes *Salmonella* antibiotic tolerance specifically through the YdiU-YdiV pathway (Fig.7c-d).

**Fig.7.**
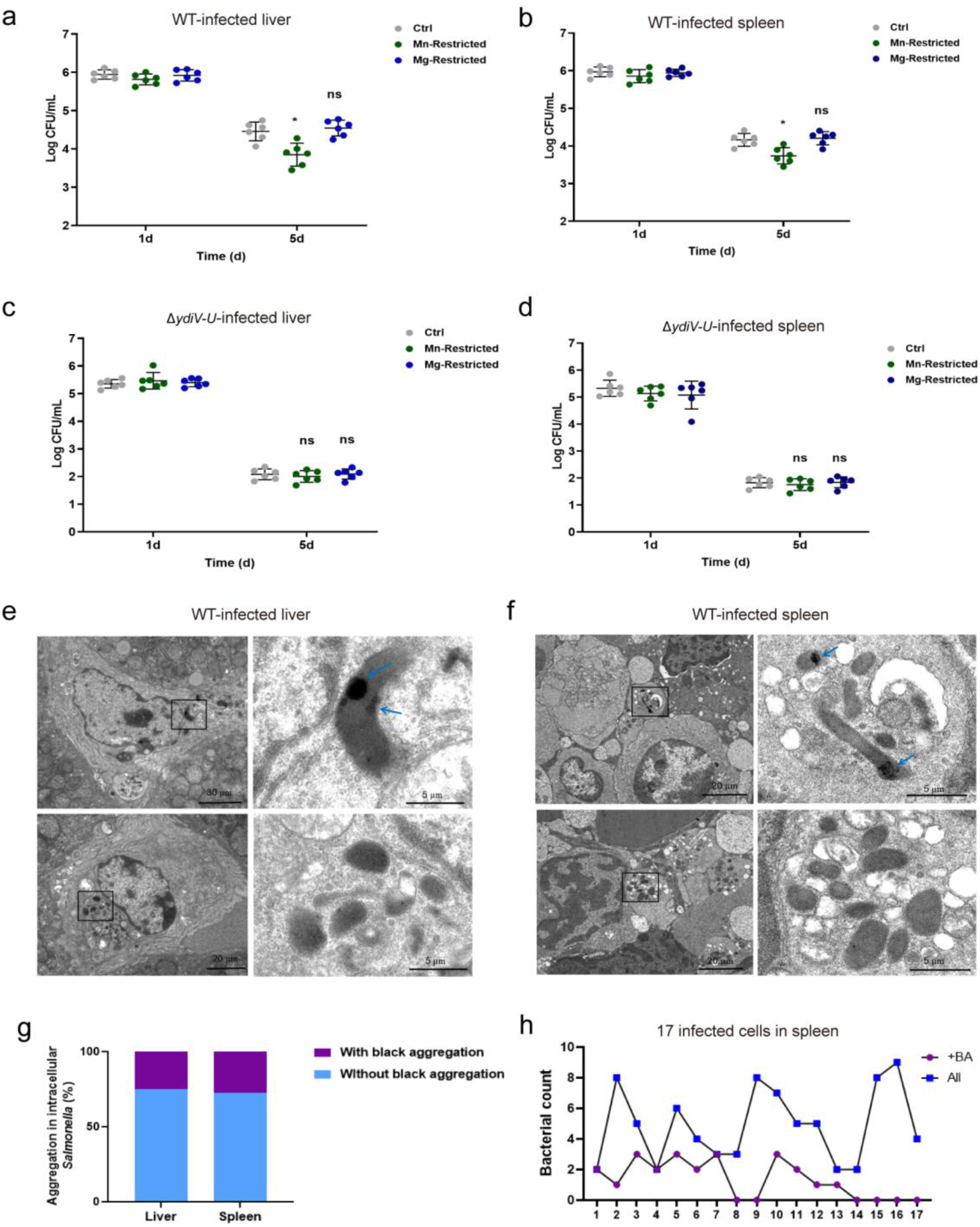
Restriction of host-derived Mn^2+^ effectively promotes *Salmonella* antibiotic clearance. (**a**-**b**) Residual bacterial counts in the liver (a) and spleen (b) of mice fed manganese-or magnesium-restricted diets, respectively, following infection with wild-type *Salmonella* and during antibiotic treatment (n = 6). (**c**-**d**) Residual bacterial counts in the liver (c) and spleen (d) of mice fed manganese-or magnesium-restricted diets, respectively, following infection with Δ*ydiV-U Salmonella* and during antibiotic treatment (n = 6). (**e**) Representative electron micrographs of bacterial ultrastructure in host liver 5 day post infection. The upper panel shows representative images of bacteria forming intracellular black aggregates, with blue arrows indicating their locations. The lower panel shows representative images of bacteria lacking intracellular black aggregates. (**f**) Representative electron micrographs of bacterial ultrastructure in host spleen 5 day post infection. (**g**) Analysis of the proportion of bacteria producing or lacking black aggregates in the livers and spleens of infected mice (Liver: n=32 bacteria; Spleen: n=83 bacteria). (**h**) Quantification of the total number of bacteria per bacteria-containing cell (All) and the number of bacteria containing black aggregates (+BA) in the spleen. Significance significance: *p < 0.05; ns, not significant.

Using transmission electron microscopy, we examined the status of intracellular protein aggregates in *Salmonella* within the livers and spleens of mice during ceftazidime treatment for wild-type *Salmonella* infection (Fig.7e-h). We observed that a significant proportion of bacteria in the infected liver (25%) and spleen (27.7%) contained intracellular aggregation foci, indicating that many pathogens entered a dormant state *in vivo*. Conversely, the absence of such aggregates in a subset of bacteria suggested their active proliferation. As the Δ*ydiV-U* strain was largely cleared after five days of treatment, we obtained insufficient bacteria within the tissues to evaluate protein aggregate formation in these bacteria. Collectively, our results demonstrate that *Salmonella* employs YdiU-YdiV-mediated protein aggregation to dynamically modulate its growth rate within the host, thereby evading antibiotic clearance.

## DISCUSSION

Emerging evidence positions antibiotic tolerance as a critical precursor to therapeutic failure through two primary facts: First, tolerant bacterial populations facilitate the emergence of resistant mutants by prolonging pathogen survival under antibiotic exposure^63–65^. Second, clinical treatment failures often arise from slowly proliferating resistant subpopulations exhibiting tolerance phenotypes, rather than the numerically limited persister cells^5,34^. This underscores the urgent need to elucidate the molecular mechanisms underlying bacterial phenotypic tolerance in host-pathogen interactions, in order to address the therapeutic challenges posed by recalcitrant infections.

Here, we elucidate a biphasic growth curve of antibiotic-tolerant *Salmonella* within macrophage, characterized by a distinct lag phase with minimal reproductive activity followed by a pronounced logarithmic (log) phase marked by rapid proliferation. We further discovered the potential mechanism (Extended Data Fig.10). First, *Salmonella* infection triggers transient upregulation of Mn²⁺ in macrophages. Meanwhile, antibiotic-induced stress triggers Mn^2+^ uptake of *Salmonella*, activating the UMPylator YdiU. YdiU-mediated UMPylation induces widespread protein aggregation, effectively arresting core cellular processes and establishing a protective dormant state. The transition from dormancy to resuscitation is governed by a Mg^2+^-dependent counterbalance system: as environmental antibiotic concentrations decline, Mg^2+^ accumulation promotes YdiV-mediated deUMPylation, restoring protein functionality and resuscitating translational activity. This metal-mediated regulation allows *Salmonella* to transiently halt growth under antibiotics while preserving revival capacity, evading both antimicrobial effects and immune clearance during macrophage infection. Our findings challenge the conventional paradigm that antibiotic-induced growth retardation primarily results from passive nutrient limitation. Instead, we demonstrate that *Salmonella* actively modulates its growth dynamics through stress-responsive regulation. This biphasic adaptation represents an evolutionarily optimized strategy rather than passive metabolic failure.

This study elucidates the mechanism underlying the widespread growth heterogeneity observed during *Salmonella in vivo* infections^34,35,37–41^. Our single-cell resolution approach uncovers two distinct phases of antibiotic-tolerant *Salmonella* during macrophage infection: an YdiU-regulated lag phase enabling antibiotic evasion, followed by YdiV-dependent exponential growth facilitating host exploitation. As a prototypical intracellular pathogen, *Salmonella*’s infection chronology within individual host cells exhibits critical temporal variations often masked by population-averaged measurements. We propose that future investigations should employ single-cell spatiotemporal tracking to resolve infection-stage-specific bacterial adaptations, as asynchronous invasion timing fundamentally confounds bulk experimental readouts in macrophage infection models.

Previous studies have found that protein aggregation is associated with bacterial dormancy and persistence, with the extent of aggregation exhibiting a positive correlation with the depth of dormant states^20–23^. Furthermore, a dormant-specific subcellular structure, termed the regrowth-delay body (RDB), emerges as a critical regulator orchestrating bacterial phenotypic switching between dormancy and resuscitation^59^. Here, we demonstrate that intramacrophage antibiotic-tolerant *Salmonella* harbor RDB-like protein aggresome, triggering by endogenously generated UMPylation-regulated protein aggregation. Initial quantitative proteomic profiling identified 56 UMPylated peptides corresponding to 46 distinct *Salmonella* proteins upon YdiU induction^45^. Comparative analysis revealed that the majority of YdiU-mediated UMPylated substrates colocalized with dormancy-related protein aggregates, thereby providing another evidence supporting the mechanistic link between YdiU’s enzymatic activity and bacterial dormancy through UMPylation-dependent protein aggregation.

UMPylation-induced protein phase separation exhibits universality. UMPylation induces phase separation of all detected substrates *in vitro*, suggesting that the UMP moiety may be a critical molecular glue driving phase separation. The UMP moiety facilitates hydrophobic π-π interactions via its aromatic structure, while numerous studies underscore the significance of π-π interactions in the formation of LLPS^66–68^. Additionally, UMPylation preferentially targets tyrosine/histidine residues in hydrophobic/acidic sequence contexts, amplifying π-stacking capacity through aromatic side chains. Our findings provided a mechanism for protein aggregates formation and establish prokaryotic PTM-mediated phase separation as a survival strategy while providing a framework for analyzing microbial stress responses through molecular crowding dynamics. The broad substrate spectrum of YdiU suggests UMPylation coordinates proteome-wide functional reorganization rather than individual protein regulation, opening new avenues for studying bacterial tolerance mechanisms through phase behavior modulation.

As a core component of bacterial translational machinery, TufA demonstrates extraordinary abundance, accounting for 5%-10% of the total proteomic mass in actively replicating bacterial cells^69^. Extensive research has elucidated the regulatory mechanisms governing its post-translational modifications^50,53,54,70–72^. This study demonstrates that UMPylation alters the structure of TufA, thereby triggering its phase separation. In fact, TufA lacks internal disordered regions in its sequence, and there are no existing reports of its ability to undergo LLPS. This paradigm-shifting discovery raises fundamental questions about the broader implications of PTM-mediated structural remodeling in protein phase behavior, necessitating systematic interrogation of other known TufA modifications.

YdiV, encoded by a gene co-localized with *ydiU* within the same operon, was identified as a de-UMPylator in this study. YdiV is the first protein confirmed to exhibit this specific activity. Notably, YdiV belongs to the EAL-like protein family, which primarily functions to hydrolyze the signaling molecule c-di-GMP^56^. While numerous EAL mutant proteins with active-site mutations are previously classified as catalytically inactive in bacterial genome, our study suggests that these proteins may retain enzymatic activity by catalyzing alternative reactions.

The regulation of UMPylation and de-UMPylation during infection depends on host-derived metal ions. After infection, mitochondrial Mn^2+^ are released into the host cytoplasm, triggering innate immune responses and autophagic flux^73,74^. Elevated intracellular Mn^2+^ concentrations synergize with antibiotic stress to drive Mn^2+^ influx in bacteria. Mn^2+^ further activates the UMPylation machinery within bacteria. With prolonged infection, as antibiotic efficacy wanes, bacterial cells undergo Mn^2+^ efflux and concomitant Mg^2+^ accumulation, which initiates deUMPylation-mediated resuscitation. This precisely coordinated metalloregulatory axis establishes spatiotemporal control over pathogen proliferation through bimodal enzymatic modulation. Recent studies have revealed that *Salmonella*’s precise regulation of Mn^2+^ efflux pumps and transport systems within macrophages closely aligns with the dynamic Mn^2+^-dependent regulatory mechanisms across different infection stages described in this work^44^. Our study reveals two novel insights: First, Mn^2+^ not only aids bacterial antioxidant defense via SOD activation but critically enables dormancy and antibiotic tolerance. Second, despite host magnesium scarcity, pathogens acquire significant Mg^2+^ during late infection to reactivate dormant bacteria.

Our study reveals the critical role of host metal ions and the YdiV-YdiU axis in mediating *Salmonella* antibiotic tolerance. Strikingly, deletion of the YdiU-YdiV system reduced bacterial tolerance to antibiotics by up to 100,000-fold in murine liver and spleen, enabling complete host clearance of the double-knockout strain within one week. These findings identify the YdiU/YdiV pathway as a promising therapeutic target; developing inhibitors against YdiU and YdiV, particularly in combination with conventional antibiotics, could provide an effective strategy for controlling persistent *Salmonella* infections. Furthermore, we demonstrate that host Mn²⁺ are essential for *Salmonella*’s *in vivo* antibiotic tolerance. This discovery carries significant clinical implications: While Mn²⁺ is recognized as an immunological adjuvant due to its ability to activate innate immunity^73^, our data reveal that exogenous Mn²⁺ supplementation simultaneously enhances bacterial resistance to antibiotic clearance. Consequently, the safety profile of Mn²⁺-based adjuvants requires urgent reassessment in the context of bacterial co-infections.

## AUTHOR CONTRIBUTIONS

B.L. and Q.C. designed the study; W.W., T.S. and X.Y. performed experiments concerning phase separation; W.W., Q.B. and Y.Z. performed electron microscopy and real-time imaging. R.L., X.L. and Z.M. performed *in vitro* experiments of persister formation; H.J., R.X., and N.S. performed experiments related to YdiV and de-UMPylation; YY.W., Y.Y., YZ.W. and X.Z. performed *In vivo* experiments on cell and animal infections; B.L., Q.C., W.W., R.L., T.S., H.J., and YY.W. analyzed and interpreted data; B.L. wrote the manuscript with input from the other authors.

## ACKNOWLEDGEMENT

We acknowledge Dr. Wenyi Du (Sichuan Model Technology Co., Ltd.) for his help in molecular simulations. This work was supported by the National Natural Science Foundation of China [32170034, 22107059, 81902038 and 31900124], the Taishan Scholar Project of Shandong Province [tsqn202211216 and tsqn202211221], the Natural Science Foundation of Shandong Province [ZR2023YQ060, ZR2022YQ66, ZR2024MC045, ZR2024QC015 and ZR2024QC209], the Youth Innovation Technology Program innovation team of Shandong Provincial University [2023KJ170], the Joint Innovation Team for Clinical & Basic Research of Shandong First Medical University [202410]. the Key Project of Medical and Health Science and Technology in Shandong Province [202401060550].

## METHODS

### Bacterial strains and propagation

*S.Typhimurium* ATCC 14028 and its derived strains were employed for functional studies. The culture was incubated overnight and subsequently subcultured into the corresponding medium. Once the optical density at 600 nm (OD600) reached 0.4-0.5, bacterial cells were harvested for follow-up observations or experiments. Luria-Bertani (LB) medium and M9 minimal medium were prepared as described previously^75^. H_2_O_2_-stress medium (H_2_O_2_) was LB medium supplemented with H_2_O_2_ to a final concentration of 1 mM. Acid-stress medium (H^+^) was LB medium supplemented with 0.5 μl/mL acetic acid (∼pH 5.0). Proteins were expressed in *E. coli* BL21(DE3). The culture was grown overnight and then subcultured into LB medium containing the appropriate antibiotics. When the optical density at 600 nm (OD600) reached 0.4-0.6, the cultures were cooled to 16°C and induced overnight with the addition of IPTG.

### Cell lines and culture conditions

RAW264.7 macrophage cells were cultured in Dulbecco’s Modified Eagle Medium (DMEM, Gibco) supplemented with 10% fetal bovine serum (FBS) at 37 °C in a 5% (v/v) CO_2_ atmosphere.

### Construction of *Salmonella* strains

Strains used in this study are listed in Supplementary Table S1. The Δ*ydiV-U*, Δ*corA*, Δ*mgtA*, Δ*mgtB*, Δ*zntB*, Δ*mntH* and Δ*mntP* strains were created utilizing the lambda Red recombinase system, as previously outlined^76^. In summary, the chloramphenicol resistance-FRT cassette was amplified from pKD3 or pKD4, including sequences that flank the target gene. The resulting product was introduced into the *S. typhimurium* ATCC14028 strain carrying the pKD46 plasmid, from which recombinant colonies were isolated on LB agar plates supplemented with 25 μg/ml chloramphenicol. To confirm the successful deletion of the target gene, PCR was conducted using primers positioned outside the gene of interest. The pCP20 plasmid was later applied to eliminate the chloramphenicol resistance gene. The Δ*mntP*&*fieF* double mutant strain was constructed from the Δ*mntP* parental strain.

A combination of the CRISPR/Cas9 and Red recombination systems was used to edit the genome of the strain ATCC14028S, resulting in a modified strain with an in situ insertion of GFP upstream of the TufA gene^77^. The strain editing was performed by Ubigene (Guangzhou, China). First, the Tuf-1-gRNA vector and the donor vector carrying GFP and the recombinant homologous arms were constructed. Then, both vectors were electroporated into the strain ATCC14028S using the following parameters: 1.8 kV, 5 ms, and 1 pulse. After recovery, an appropriate amount of the bacterial culture was spread onto LB plates with the corresponding antibiotics and incubated for 1-2 days. Single colonies were picked and identified by PCR using the primers Test-tuf-1-F: GAACGCTATCGGCATAGGCT and test-tuf-1-R: ATCCCGTGCTCTCTCCTGAA. Positive clones were selected for sequencing and preservation. The YdiV E116A mutant strain was performed by Ubigene (Guangzhou, China).

### Plasmid construction

Primers and plasmids used in this study are listed in Supplementary Table S2 and Table S3. For the construction of the TufA expression plasmid, DNA fragments of TufA of *E. coli* strain K-12 substrain MG1655 was cloned into the pGL01 vector, a modified pET15b vector containing an PPase cleavage site to remove the His tag. For the in vivo study, the *Salmonella tufA* gene was amplified and cloned into the pFPV25.1 vector. The resulting recombinant plasmid, pFPV25.1-TufA, facilitated the expression of a GFP-fusion TufA protein. To construct the GFP-fusion plasmids, DNA fragments of *tufA* and other genes (*fur*, *flhD* and *groEL*) were fused to the GFP gene using a flexible linker sequence (GSGSGSGSGSGS). For the deUMPylation assay, the full-length YdiV protein was amplified from the genomic DNA of *Salmonella Typhimurium strain 14028S* and cloned into the expression vector pGL01. Mutants of YdiV (E29A, E116A, D165A, C193A) were generated using a two-step PCR strategy and individually cloned into pGL01. YdiU and TufA genes were obtained by PCR amplification from *Salmonella typhimurium* ATCC 14028s genomic DNA and cloned into the pFPV25.1 vector containing a GFP report gene for *in vivo* study.

### Protein expression and purification

The recombinant plasmids were transformed into *E. coli* strain BL21 (DE3) for protein expression. For YdiV and its mutants, the recombinant *E. coli* cells were induced with 0.1 mM isopropyl β-D-1-thiogalactopyranoside (IPTG) when the optical density at 600 nm (OD600) reached 0.4–0.6, followed by incubation at 37°C for 3 hours. For other proteins, the *E. coli* cells were cultured to an OD600 of 0.4–0.6, then cooled to 16°C and induced overnight with 0.01–0.1 mM IPTG. After induction, the cells were collected, lysed, and the proteins were purified using a Ni²⁺-NTA affinity column. The His-tags were subsequently removed by PPase treatment. Finally, the proteins were concentrated and further purified using Superdex 200 chromatography. Notably, the PPase treatment step was omitted during the purification of the GFP fusion protein.

### *In vitro* antibiotic resistant curves

The *Salmonella* strains were cultured overnight in standard liquid LB medium at 37°C. The cultures were then diluted 1:100 in fresh minimal M9 medium and allowed to grow until an optical density at 600 nm (OD600) of 0.3 was reached. Subsequently, antibiotics were added: gentamicin at 50 µg/mL, norfloxacin at 4 µg/mL, or ceftazidime at 32 μg/mL. Samples were taken at 2, 4, 6, and 24 hours, where 1 mL of bacterial cells was collected by centrifugation (5000 rpm for 5 minutes) and washed twice with PBS to remove the antibiotics. The cells were then resuspended in 1 mL of PBS. A total of 100 µL of this suspension was plated on LB agar plates, and CFUs were counted to determine the number of viable bacteria.

### Antibiotic tolerance of *Salmonella* within macrophage

For the data shown in Fig.1c-e, Fig.4o-p, Fig.5g and Fig.6l-m, RAW264.7 cells were seeded in 24-well plates at 5×10⁵ cells/well. Single colonies of relevant bacterial strains were inoculated into standard LB medium and cultured overnight at 37°C with 200 rpm shaking. Activated strains were subcultured in high-salt LB medium to OD₆₀₀ ≈ 0.5. Bacterial cells were pelleted by centrifugation (3,000-4,000 × g, 5 min), washed, and resuspended in DMEM to 1×10⁸ CFU/mL. Cells were infected at MOI=10 for 1 h. For initial bacterial load (0 h), infected cells were washed thrice with PBS, lysed with 1% Triton X-100, serially diluted, and plated. For time-course analysis, infected cells received antibiotic-containing medium (ceftazidime 32 μg/mL or norfloxacin 4 μg/mL or PBS as control) and were sampled at indicated time points (0, 6, 12, 18, 24 h). Colony counting was performed post-serial dilution. All procedures were conducted in a biosafety cabinet with incubation at 37°C/5% CO₂.

For the data shown in Fig.6i, RAW264.7 cells were seeded in 24-well plates at 5×10⁵ cells/well while concurrently activating wild-type *Salmonella*; the following day, bacterial cultures were subcultured 1:100 into high-salt LB medium and incubated at 37°C to OD₆₀₀ ≈ 0.5, followed by centrifugation, washing, and resuspension in DMEM to 1×10⁸ CFU/mL. Prior to infection at MOI=10, cell medium was replaced with DMEM containing either 50 μM metal ions (MnCl₂, MgCl₂, ZnCl₂, CaCl₂), 0.4 mM EDTA, or equivalent-volume H₂O₂ control. After 1 h infection, initial bacterial load (0 h) was determined by PBS washing, 1% Triton X-100 lysis, and serial 10-fold dilution plating; infected cells then received complete medium with corresponding compounds plus antibiotics (ceftazidime 32 μg/mL or norfloxacin 4 μg/mL), with intracellular bacteria quantified identically after 18 h incubation. All procedures were performed in a biosafety cabinet at 37°C/5% CO₂.

### Antibiotic tolerance of *Salmonella* in animal tissues

All animal experiments were approved by the Animal Care and Use Ethics Committee of the Institute of Basic Medical Sciences, Shandong Academy of Medical Sciences (IBMSAMSC No.097). The subjects of the experiment were BALB/c mice, each weighing between 16 and 18 g, which were randomly divided into groups. Prior to infection, all mice were administered antibiotics (ceftazidime at a dosage of 0.06 mg), beginning in the afternoon before infection and followed by subsequent injections every 12 hours. Bacterial strains were then injected intraperitoneally into each group of mice at the indicated concentration of 2×10^6^ CFU/mouse. In Fig.1b, the mice were randomly re-divided into four groups (WT, WT+ceftazidime, Δ*ydiU*, and Δ*ydiU*+ceftazidime), with five mice in each group. All mice received either antibiotic injections (ceftazidime at a dosage of 0.06 mg) or PBS injections in the afternoon prior to infection, with continued injections administered every 12 hours thereafter (24h for Fig.5i). The survival of the remaining mice was monitored and recorded throughout the duration of the experiment.

For data presented in Fig.5i and Fig.6n-o, in the indicated time post-infection (0d: 6hpi, 2d: 48hpi, 5d: 120hpi), the mice were processed according to the experimental protocol at designated time points, and their livers and spleens were harvested. The mouse livers and spleens were lysed in 1% Triton X-100 in physiological saline (PBST). Subsequently, all samples were diluted with PBST, spread onto LB agar plates, incubated at 37°C for 24 hours, and quantitatively analyzed for CFU.

For data presented in Fig.7, mice were randomly divided into four groups: normal diet, Mn-restricted diet, and Mg-restricted diet groups; after five days of dietary conditioning, wild-type *Salmonella* and Δ*ydiV-U* double-knockout strains were activated one day pre-infection, with concurrent initiation of ceftazidime pretreatment (0.06 mg/dose, i.p.). The following day, bacterial cultures were subcultured 1:100 into high-salt LB medium, grown to OD₆₀₀ ≈ 0.5, pelleted (5,000 × g, 5 min), and resuspended in PBS at 2×10^6^ CFU/mouse for intraperitoneal infection. At 6 h post-infection (day 0), subset mice were sacrificed; remaining animals continued ceftazidime treatment (q12h). The mouse livers and spleens were lysed in 1% Triton X-100 in physiological saline (PBST). Subsequently, all samples were diluted with PBST, spread onto LB agar plates, incubated at 37°C for 24 hours, and quantitatively analyzed for CFU.

### *In vitro* protein synthesis inhibition assay

An *in vitro* cell-free translation system, utilizing a luciferase reporter (Promega), was employed to assess the inhibitory effects of YdiU on protein synthesis. Specifically, the reaction mixture was prepared following the manufacturer’s instructions and consisted of *E. coli* S30 extract, S30 premix without amino acids, an amino acid mixture lacking methionine, an amino acid mixture lacking leucine and a reporter plasmid containing the luciferase gene. A final concentration of 0.1 µM YdiU was added to form the experimental group. An equal volume of SD buffer (10 mM Tris, 100 mM NaCl, pH 8.0) served as the positive control, while the sample without the luciferase plasmid acted as the negative control. After gently mixing the components, the mixture was incubated at 37°C for 2 hours. The reaction was then halted by placing the mixture in an ice bath for 5 min, after which it was used for subsequent western blot analysis and luciferase activity detection experiments.

### Western blotting

For the data shown in Fig.2b, the reaction mixture was combined with SDS-PAGE loading dye and heated at 95°C for 10 minutes before gel electrophoresis. Following separation, proteins were transferred to nitrocellulose membranes at 250 mA for 2 hours. The membranes were blocked with 5% milk in phosphate-buffered saline with 0.1% Tween (PBST) for 2 hours at room temperature, then incubated overnight at 4°C with primary antibodies against luciferase (Abcam, 1:10,000) and GapA (lab made, 1:10,000) in PBST. After washing three times with PBST, membranes were incubated for 1 hour at 25°C with HRP-conjugated secondary antibodies (Goat anti-Rabbit IgG, Abcam). Following three additional washes with PBST, proteins were visualized using a chemiluminescent substrate and detected with a FluorChem imager. Protein levels were quantified using ImageJ software.

### Luciferase Bioluminescence Assay

For the data shown in Extended Data Fig.1c, the luciferase expression was measured with its activity. After spotted into the well of a white 96-well plate, 10 µL reaction mixture was added 50 µL of room-temperature Luciferase Assay Reagent (promega) and mixed quickly by pipetting. Light emission was recorded with a luminescence counter (PerkinElmer). For the data shown in Fig.2c, the recombinant strains harboring pFPV25.1-Luc were routinely cultured in LB medium at 37°C. Bacterial cell lysis buffer was prepared by adding 20 mg of lysozyme (Shangon Biotech) to 1 mL of Lysozyme Buffer (Shangon Biotech). After 5, 15, and 20 hours, 1 mL of culture was centrifuged at 12,000 rpm for 3 minutes, washed twice with sterilized water to remove the medium, and then suspended in 100 µL of bacterial cell lysis buffer. Following a 15-minute incubation at room temperature, the mixture was centrifuged at 12,000 rpm for 3 minutes. Subsequently, 100 µL of the supernatant was combined with 100 µL of reaction solution, following the instructions of the Firefly Luciferase Reporter Gene Assay Kit (Beyotime), and the luminescence (relative light units, RLU) was measured.

### *In vitro* UMPylation assay

For the data shown in Fig.2d-e, the biotin-labeled UTP (biotin-16-UTP) was used as the UMP donor. A 20 µL reaction buffer consisting of 25 mM Tris-HCl (pH 7.5), 10 mM dithiothreitol (DTT), 25 mM NaCl, 0.625 mM MnCl_2_, and 50 mM biotin-16-UTP was prepared, to which 2 µL of *E. coli* S30 extract from an *in vitro* translation system (Promega) was added with or without 2 μg of YdiU. The mixture was incubated at 30°C for 1 hour. Following incubation, the reaction was combined with SDS-PAGE loading dye and heated at 95°C for 10 minutes. Subsequently, a 12.5% SDS-PAGE and streptavidin-HRP blot were conducted as previously described^45^. For the experiment shown in Fig.2g, Extended Data Fig.4c-d, Cy3-UTP or Cy5-UTP was utilized as the UMP donor. Purified proteins (5 μg of TufA or GFP-TufA) were incubated with or without 2 μg of YdiU in a 20 µL reaction buffer comprising 25 mM Tris-HCl (pH 7.5), 10 mM dithiothreitol (DTT), 25 mM NaCl, 0.625 mM MnCl_2_, and 25 µM Cy3-UTP or Cy5-UTP. After incubation at 30°C for 1 hour, the reaction mixture was analyzed via SDS-PAGE or visualized using microscopy. For the *in vitro* UMPylation assay related to the data shown in Fig.2h-i, Extended Data Fig.3a-f, the same method was employed, except that UTP substituted Cy3-UTP or Cy5-UTP in the reaction.

### Biotin-streptavidin pull down assay

For the data shown in Fig.2e, a biotin-streptavidin pull-down assay was utilized to identify the UMPylated target proteins from *E. coli* S30 extract catalyzed by YdiU. Following three washes with SD buffer (10 mM Tris, 100 mM NaCl, pH 8.0), 50 µL of streptavidin beads (Thermo Scientific) were incubated with 200 µL of the UMPylated sample for 30 minutes at 25°C. The beads were then collected by centrifugation and subjected to three additional washes with SD buffer. Finally, the target proteins bound to the beads were eluted using SD buffer containing 8 M urea. Then, the elution was analyzed by SDS-PAGE and subsequent mass spectrometry identification.

### Mass spectrometry

For the data presented in Fig.2e and Extended Data Fig.6, samples were separated on a 12.5% SDS-PAGE gel and stained with Coomassie Brilliant Blue. The stained bands were excised and stored at 4°C. In-gel digestion, LC-MS/MS analysis, and data processing were conducted by Applied Protein Technology Co. Ltd. In brief, the excised gel pieces underwent reduction, alkylation, and were then digested with trypsin at 37°C for 20 hours. The resulting digestion products were desalted, dried, and resuspended in 0.1% formic acid solution. The tryptic peptides were subsequently loaded onto a trap column and analyzed using a Q Exactive mass spectrometer (Thermo Scientific) coupled with an EasynLC system (Thermo Fisher Scientific). Data acquisition targeted the 20 most intense peaks from each MS full scan. MS/MS spectra were automatically searched against the Uniprot *Escherichia coli* database (1856608, version 20200217) using MASCOT 2.2 software, resulting in the identification of protein fragments. The data shown in Fig.2h-i, the tryptic digestion peptides were resuspended in 2% acetonitrile/0.1% formic acid solution and then separated with an EASY-nLC 1200 UPLC system. The peptides were subjected to NSI source followed by tandem mass spectrometry (MS/MS) in Q Exactive HF-X (Thermo Fisher) coupled online to the UPLC. The resulting MS/MS data were processed using Proteome Discoverer 2.4 software. Tandem mass spectra were searched against *Escherichia coli* (strain K-12) database (UniProt no. 83333) database. UMPylated peptides were identified by searching for the “PhosphoUridine” modification.

### Bacterial Two-Hybrid assay

The *tufA* and *ydiU* genes were amplified from the genomic DNA of *Salmonella typhimurium* ATCC 14028s using polymerase chain reaction (PCR). Plasmids pUT18C-ydiU and pKNT25-tufA were constructed and subsequently co-transformed into *E. coli* BTH101 via electroporation. Control strains were created by transforming BTH101 with plasmids pKNT25 and pUT18C. The strains were cultivated to the logarithmic growth phase in LB medium supplemented with 100 µg/mL ampicillin, 50 µg/mL kanamycin, and 0.5 mM IPTG. Subsequently, 2 µL of each sample was plated on LB-X-Gal plates containing 100 mg/mL ampicillin, 50 mg/mL kanamycin, 0.5 mM IPTG, and 40 mg/mL X-Gal. β-galactosidase activity was assessed using a β-galactosidase Activity Assay Kit (Solarbio), following the manufacturer’s instructions.

### *In vitro* phase separation assay

To prepare UMPylated proteins, purified protein was incubated with or without 1µM YdiU in a reaction buffer containing 25 mM Tris-HCl (pH 7.5), 10 mM dithiothreitol (DTT), 25 mM NaCl, 0.625 mM MnCl₂, and 50 µM UTP. Following a 1-hour incubation at 30°C, the reaction mixture was desalted and diluted in a phase separation buffer. For GFP-TufA, the phase separation solution consisted of 25 mM Tris (pH 7.6), 25 mM NaCl, 10 mM DTT, 10 µM GTP, 0.25 µg/µL tRNA, and 2% PEG 2000. In contrast, the phase separation solution for the other GFP fusion proteins (Fur, FlhD, GroEL) contained 25 mM Tris (pH 7.6), 25 mM NaCl, 10 mM DTT, and either 2% PEG 2000 or 1% PEG 3000. Then, the sample was loaded into a confocal dish and imaged using a ZEISS LSM 980 Confocal Microscope equipped with a ×40 objective.

### *In vivo* phase separation assay

For the data presented in Extended Data Fig.3g-h, activated bacterial cells were cultivated in M9 medium until reaching the stationary phase. Fresh cultures were then diluted 10-fold in PBS buffer and mixed with an equal volume of a 50% agar-water solution, which had been cooled to approximately 50°C. Subsequently, 10 μL of the resulting mixture was placed on a confocal dish for imaging with a confocal microscope. For the data presented in Fig.3d-g and 4k-l, bacterial cells were cultivated overnight and then subcultured into the appropriate medium until they reached the stationary phase. The oxidative stress condition was achieved using LB medium supplemented with H_2_O_2_ to a final concentration of 1 mM, while the acidic stress medium consisted of LB supplemented with 0.5 µL/mL acetic acid, adjusted to a pH of 5.0. The subsequent steps were the same as described above.

### Fluorescence microscopy

Samples for Fig.3, Fig.4 and Extended Data Fig.3 were observed with a LSM 980 (Zeiss) inverted microscope, 63x oil lens and DIC filter. Image capture and reconstruction of high resolution SIM2 images were performed with Elyra 7 (Zeiss). All images were analysed using Zeiss ZEN Blue or ImageJ softwares.

### Fluorescence Recovery After Photobleaching (FRAP)

The *in vitro* phase separation assays were reconstituted as described above and samples were observed immediately after mixing using an inverted laser scanning confocal microscope (LSM 980, Zeiss), and pulsed supercontinuum white light source set at emission wavelength 488 nm. Droplets set at the bottom of the coverslip were point bleached for 100 ms with a laser power at 8.0% achieving a 50-60% fluorescence reduction at the bleached spot. A single image was acquired immediately before and one immediately after bleaching and then every 5 sec to reduce photobleaching from image acquisition. Image analysis was performed using ImageJ. To calculate the fluorescence recovery, the average fluorescence intensity of the bleached area over time was normalized to the fluorescence of the whole droplet.

### Electron microscopy

For the data shown in Fig.3i, 4m, 6p, Extended Data Fig.5f and Extended Data Fig.8a, RAW264.7 cells infected with *Salmonella* at a multiplicity of infection (MOI) of 10 over various durations involved several steps. At designated time points, the cell culture medium was replaced with an electron microscopy fixation solution, and cells were fixed at room temperature in the dark for 5 min. For the data shown in Extended Data Fig.5d and 6a, activated bacterial cells were grown in LB medium with 0.1% L-arabinose to stationary phase (5d) or OD 0.5 (6a). The cultures were then gently centrifuged and resuspended in 500 μL of electron microscope-fixing solution. Following fixation, the cells were collected and resuspended in fresh fixation solution, undergoing an additional 30-minute fixation at room temperature before being stored at 4°C. A 1% agarose solution was used for pre-embedding, followed by fixation in 1% osmium tetroxide at room temperature for 2 hours. Samples were washed twice with PBS, dehydrated with ethanol and acetone, then embedded and dried overnight at 37°C. After polymerization at 60°C for 48 hours, thin sections were obtained using a Leica UC7 ultramicrotome, stained, and mounted on a 150-mesh copper grid. The grid was air-dried overnight in the dark before imaging with a Hitachi HT7800/HT7700 transmission electron microscope. The electron microscopy analyses were carried out with support from Servicebio Co., Ltd.

For the data shown in Fig.7, sterile spleen and liver tissues were collected and immediately preserved in pre-chilled (4 °C) electron microscopy fixative. Following thorough rinsing with 0.1M phosphate buffer (PB, pH 7.4), the tissue samples were fixed in 1% osmium tetroxide solution in the dark for 2 hours. Subsequent processing included dehydration through a graded ethanol series (30%-100%) and acetone, followed by infiltration and embedding in Epon812 resin. Polymerization was carried out at 60°C for 48 hours. Semi-thin sections (1.5 μm) were stained with toluidine blue for light microscopic localization. Ultrathin sections (60-80 nm) were double-stained with uranyl acetate and lead citrate before being examined under a transmission electron microscope (TEM) for image acquisition. All experimental procedures were strictly performed within a Class II biosafety cabinet to ensure compliance with biosafety requirements. The grid was air-dried overnight in the dark before imaging with a Hitachi HT7800/HT7700 transmission electron microscope.

### *In vitro* de-UMPylation assay

To prepare self-UMPylated YdiU, 5 μg of purified YdiU^475^ was incubated with 10 μL of reaction buffer containing 25 mM Tris-HCl pH 7.5, 1 mM DTT, 100 mM NaCl, 10 mM MnCl₂, and 500 μM Cy5-UTP at 30°C for 60 min. To prepare UMPylated TufA, 6 μg of TufA were incubated with 1 μg of YdiU^475^ in a 10 μL reaction buffer. Prior to de-UMPylation, the UMPylation products were desalted using a desalting column (Cytiva) and eluted in SD buffer (10 mM Tris-HCl, pH 8.0, and 100 mM NaCl). The de-UMPylation reactions were conducted using 1 μg of UMPylated YdiU (UMP-YdiU) or UMPylated TufA (UMP-TufA), alongside varying concentrations of YdiV or its mutants, in a 10 μL reaction buffer (containing 10 mM Tris-HCl, pH 8.0, 100 mM NaCl, 20 mM CaCl_2_, and 1 mM DTT) at 30°C for designated time points. The samples were subsequently analyzed using a 12% NuPAGE gel and visualized via autoradiography. An *in vitro* depolymerization assay mediated by YdiV was performed using the UMPylated GFP-TufA protein (catalyzed by YdiU and prepared as described above). After desalting, the protein was incubated with approximately 1 μM of either wild-type YdiV or its mutant variants, while a protein-free buffer served as the control. Following a 1-hour reaction period, the aggregation status of multimers was assessed.

### Circular dichroism (CD)

The protein samples were desalted and diluted in 10 mM phosphate buffer (0.73 mM Na_2_HPO_4_, 0.45 mM NaH_2_PO_4_, pH 7.2) to final concentrations of 50 or 75 µg/mL. The CD spectra were then recorded using a JASCO J-810 spectropolarimeter in a 1.0 cm quartz cell, with measurements taken between 190 nm and 250 nm at 25°C.

### Dynamic light scattering (DLS)

For the data shown in Extended Data Fig.3d-f, dynamic light scattering (DLS) was used to determine the hydrodynamic radius (rH) of protein samples. The samples were diluted to 0.5 mg/mL in SD buffer and loaded into a 96-well plate, with 100 µL of sample per well. Measurements were performed at 25.0 ± 0.1℃ using a DynaPro plate reader 3 (Wyatt Technology). Data analysis was automatically handled by the instrument’s accompanying software.

### Growth curves

For the data shown in Extended Data Fig.1e, the growth curves of the Δ*ydiU*-pBAD24::YdiU strain were assessed using the Bioscreen C Automatic Growth Analyzer (Thermo Electron Corporation). Fresh cultures were diluted 100-fold in M9 medium and transferred to the wells of a honeycomb microplate, resulting in a final volume of 300 μL. The absorbance at 600 nm was recorded every 5 minutes at 37°C over a 24-hour period. L-arabinose was added at a concentration of 0.1% when the OD600 values reached 0.3, 0.5, and 0.7, respectively.

### Molecular docking of YdiV and UMP-Tyr

Molecular docking was performed using Autodock software. The crystal structure of the YdiV (PDB code: 3TLQ) served as the receptor^58^. The three-dimensional structure of UMP-Tyr was constructed, and the MOPAC program was utilized to optimize the ligand structure and calculate the PM3 atomic charges^78^. Following this, Autodock Tools 1.5.6 was employed to prepare the receptor and ligand structures for docking^79^. The docking box was designed to encompass the entire protein, with center coordinates set at (-5.9, -3.6, -17.5). The grid dimensions were specified as 60×60×60 in the XYZ directions, with a grid spacing of 0.375 Å. A total of 200 docking runs were conducted, while all other parameters were maintained at their default settings. Energy optimization was performed using the Amber14SB force field^80^, executed in two steps: initially, a steepest descent method was applied for 2000 steps, followed by further refinement using a conjugate gradient method for an additional 2000 steps. The final optimized structures served as models for subsequent analysis.

### Molecular dynamics simulations of YdiV and UMP-Tyr

Molecular dynamics (MD) simulations were conducted using Gromacs 2021.5 software under conditions of constant temperature and pressure with periodic boundary conditions^81^. The Amber14SB all-atom force field was applied along with the TIP3P water model. During the MD simulations, LINCS algorithm was employed to constrain all hydrogen bonds, with an integration time step of 2 fs. Electrostatic interactions were calculated using the Particle-Mesh Ewald (PME) method^82^. The cutoff for non-bonded interactions was set to 10 Å, updated every 10 steps. The V-rescale method was utilized for temperature coupling, maintaining the simulation temperature at 300 K, while the Parrinello-Rahman method was used to control the pressure at 1 bar. Initially, steepest descent minimization was performed on both systems to eliminate any close atomic contacts. Subsequently, 100 ps of NVT equilibrium simulations were carried out at 300 K. Finally, six different systems underwent 50 ns of MD simulations, with conformations saved every 10 ps. The resulting simulations were visualized using Gromacs built-in tools and VMD.

### Metal ion probe incubation and imaging

For the data shown in Fig.6, RAW264.7 were cultured at 37°C with 5% CO_2_ and seeded at a density of 2 × 10^6^ per well in a 35 mm confocal dish. At various time points following *Salmonella* infection of macrophages, the culture medium was discarded, and 1 mL DMEM containing metal ion probe ( Mg^2+^: 2.5 µM, Zn^2+^: 10 µM, Ca^2+^: 4 µM) and 0.02% Pluronic F127 was added. The cells were then incubated at 37°C for 1 h. Following this, the supernatant was removed, and the cells were washed three times with pre-warmed PBS. After added 1 mL fresh DMEM medium, the cells were incubated at 37°C for an additional 30 minutes. Subsequently, 1 mL 4% paraformaldehyde was added to fix the cells at room temperature for 10 minutes. After discarding the paraformaldehyde and allowing the cells to air-dry, 2 mL of PBS was added. The samples were wrapped in foil and stored at 4°C until imaging.

The prepared samples were observed using a laser scanning confocal microscope. Imaging was performed using ZEN software, and the data were analyzed with ImageJ software. The excitation wavelengths and emission detection ranges of the fluorescent dyes used in this study are as follows: Lipi Blue channel: Ex = 405 nm, Em = 417–476 nm; MTG channel: Ex = 488 nm, Em = 500–550 nm; LDLRC channel: Ex = 561 nm, Em = 600–640 nm; Lipi Red channel: Ex = 561 nm, Em = 600–640 nm.

### Single-cell fluorescence imaging and analysis in macrophages

In the experiments depicted in Fig.5, the corresponding bacterial strains were inoculated into liquid LB medium and cultured until the OD reached 0.5. The bacterial solution was then diluted to a concentration of 5×10^5^ cells/mL and the diluted bacterial suspension was used for subsequent Raw264.7 cell invasion. Concurrently, Raw264.7 cells were cultured in DMEM medium supplemented with 10% fetal calf serum and seeded into 96-well plates at a density of 50,000 cells per well. The strains of Salmonella were subsequently introduced at a multiplicity of infection (MOI) of 10:1. After 1 hour of incubation, the cells were washed twice with PBS, and fresh DMEM medium containing 100 μg/mL gentamicin was added to eliminate any extracellular bacteria. After an additional 1 hours, the medium was replaced with DMEM containing the indicated concentrations of ceftazidime (12.5MIC: 10 µg/ml; 25MIC: 20 µg/ml; 50MIC: 40µg/ml; 100MIC: 80µg/ml) and cultured in a 37°C mini-incubator with 5% CO_2_ of THUNDER imager 3D live cell (Leica, Germany). The *Salmonella* in infected macrophages were imaged with the THUNDER imager 3D live cell Analysis Platform. Phase and fluorescence images were collected every 2 hour per well using a ×40 objective.

### Inductively coupled plasma mass spectrometry (ICP-MS)

In the experiments depicted in Fig.6b, WT *Salmonella* infected Raw264.7 cells at a multiplicity of infection (MOI) of 10:1. At various time points following *Salmonella* infection of macrophages, the culture medium was discarded, and 2 mL PBS was added. The cells were repeatedly pipetted with PBS to resuspend them in it. Following this, the samples were subjected to five cycles of freeze-thaw in liquid nitrogen for cell lysis. The supernatant was obtained after centrifugation at 12,000 rpm for 10 min at 4°C. Finally, the Mn²⁺ concentration in the supernatant was measured/determined using ICP-MS. For the data shown in Fig.6e-h, Wild-type *Salmonella* cultures were grown to an OD_600_ of 0.5, after which bacterial cells were pelleted by centrifugation and equally divided into three treatment groups: (1) the antibiotic-treated group exposed to antibiotic for 1 hour; (2) the antibiotic-removal group subjected to antibiotic treatment for 1 hour followed by centrifugation for antibiotic removal and resuspension in PBS for an additional 1 hour; and (3) the control group incubated with PBS alone for 1 hour. Above samples (0.2 ml) were introduced into Teflon digestion vessels containing 5 mL of nitric acid. Following complete reaction, the vessels were securely sealed and subjected to microwave-assisted digestion. Upon cooling to <50°C, the digested samples were carefully transferred to a fume hood for unsealing. The resulting material was reconstituted in ultrapure water and quantitatively transferred to 50-mL volumetric flasks through 3-4 cycles of rinsing. After dilution with 20 mL deionized water and filtration, elemental analysis was conducted via inductively coupled plasma mass spectrometry (Thermo iCAPQ ICP-MS). Each sample preparation included triplicate biological replicates.

### The cellular dispersion analysis of TufA

The *in vivo* phase separation of TufA was quantified using ImageJ: manual line-drawing with plot profile analysis revealed fluorescent signal aggregation at cell poles, creating intensity peaks. Normal cells showed uniform distribution. Phase separation degree was assessed by calculating peak-to-valley intensity differences along the line, enabling cross-experiment comparisons. Higher differences indicated stronger phase separation.

### Phylogenetic analysis

Full-length of YdiV amino acid sequences obtained from NCBI database were used to generate phylogenetic trees. Homologous regions were aligned using MUSCLE (https://www.ebi.ac.uk/Tools/msa/muscle/), and the alignments were adjusted manually using the BioEdit sequence alignment editor. Phylogenetic trees were generated using maximum-likelihood methods with bootstrap analysis (n=1000) using FastTree software. Trees were drawn and modified by iTOL (https://itol.embl.de/).

### Clustering analysis of aggregated proteins

Aggregated proteins was isolated and identified by mass spectrometry. Proteins with peptide-spectrum matches (PSM) exceeding 10 were selected as high-confidence identifications. Subsequent hierarchical clustering and ranking were conducted based on protein abundance (quantified by PSM values), with visualization achieved using the heatmap module in ChiPlot (https://www.chiplot.online/index.html).

### Statistical analysis

All experiments were performed three times unless otherwise stated. A two-tailed Student’s t test was used to calculate the P value using GraphPad Prism or SPSS. Error bars represent the standard deviation (SD). The statistical significance is indicated by ****P < 0.0001, ***P < 0.001, **P < 0.005, *P < 0.05.

## SUPPLEMENTAL INFORMATION

Extended Data Fig.1. YdiU inhibits protein translation and bacterial growth.

Extended Data Fig.2. Analysis of UMPylated sites on TufA.

Extended Data Fig.3. UMPylation-driven changes in the secondary structure and granularity and intracellular distribution of TufA.

Extended Data Fig.4. UMPylation-driven in vitro phase separation of TufA.

Extended Data Fig.5. UMPylation-driven phase separation of other targets and within macrophage.

Extended Data Fig.6. High conservation of deUMPylation-active residues in YdiV homologs.

Extended Data Fig.7. YdiV catalyzes de-UMPylation of YdiU.

Extended Data Fig.8. YdiV modulates protein disaggregation during infection.

Extended Data Fig.9. Proteomic identification of YdiU-mediated aggregates.

Extended Data Fig.10.Schematic diagram of *Salmonella* intracellular dormancy and resuscitation mechanism.

Table S1. Strains used in this study.

Table S2. Primers used in this study.

Table S3. Plasmids used in this study.

## Data and Code Availability

Plasmids and strains generated in this study are available from the Lead Contact. Mass spectrometry-based data were submitted to ProteomeXchange under accession number PXD057156 and PXD057051.

Original images used for the figures are deposited in the Mendeley database (http://doi.org/10.17632/x2dnvfwrdk.2).

